# The structure, catalytic mechanism, and inhibitor identification of phosphatidylinositol remodeling MBOAT7

**DOI:** 10.1101/2022.09.15.508141

**Authors:** Kun Wang, Chia-Wei Lee, Xuewu Sui, Siyoung Kim, Shuhui Wang, Aidan B Higgs, Aaron J Baublis, Gregory A Voth, Maofu Liao, Tobias C Walther, Robert V Farese

## Abstract

Cells remodel glycerophospholipid acyl chains via the Lands cycle to adjust membrane properties. Membrane-bound *O*-acyltransferase (MBOAT) 7 acylates lyso-phosphatidylinositol (lyso-PI) with arachidonyl-CoA. *MBOAT7* mutations cause brain developmental disorders, and reduced expression is linked to fatty liver disease. Further, increased *MBOAT7* expression is linked to hepatocellular and renal cancers. The mechanistic basis of MBOAT7 catalysis and substrate selectivity are unknown. Here, we report the structure and a model for the catalytic mechanism of human MBOAT7. Arachidonyl-CoA and lyso-PI access the catalytic center through a twisted tunnel from the cytosol and lumenal sides, respectively. N-Terminal residues on the ER lumenal side determine phospholipid headgroup selectivity: swapping them between MBOATs 1, 5, and 7 converts enzyme specificity for different lyso-phospholipids. Finally, the MBOAT7 structure and virtual screening enabled identification of small-molecule inhibitors that may serve as lead compounds for pharmacologic development.

## Introduction

Phospholipids are synthesized in the endoplasmic reticulum (ER) via a series of enzymatic steps that sequentially add acyl chains to a glycerol backbone^1,2^. Glycerol acyltransferase enzymes^3^ preferentially add a saturated fatty acid to the *sn-1* position, and acylglycerol acyltransferase enzymes usually esterify an unsaturated fatty acid to the *sn*-2 position. To adjust membrane properties according to need and to store specific fatty acids as precursors for bioactive lipids, cells remodel the phospholipid acyl chains in enzymatic reactions known as the Lands cycle^4^. In this process, the *sn*-2 fatty acyl chain is removed by hydrolysis, and the lysophospholipid product is re-esterified with an acyl-CoA to form a phospholipid with a different acyl chain^4,5^.

Re-acylation is carried out by lysophospholipid acyltransferase enzymes, including various membrane-bound *O*-acyltransferases (MBOAT), such as MBOATs 1, 2, 5, and 7 in humans^5^. MBOAT1 and MBOAT5 remodel phosphatidylserine (PS) and phosphatidylcholine (PC), respectively^6,7^. MBOAT2 remodels phosphatidylethanolamine (PE) and phosphatidic acid (PA), and to a lesser extent PC and PS^7^. MBOAT7 uniquely remodels phosphatidylinositol (PI), catalyzing preferentially the esterification of arachidonyl CoA to lyso-PI^5,7^ (Fig. 1a).

**Fig. 1.**
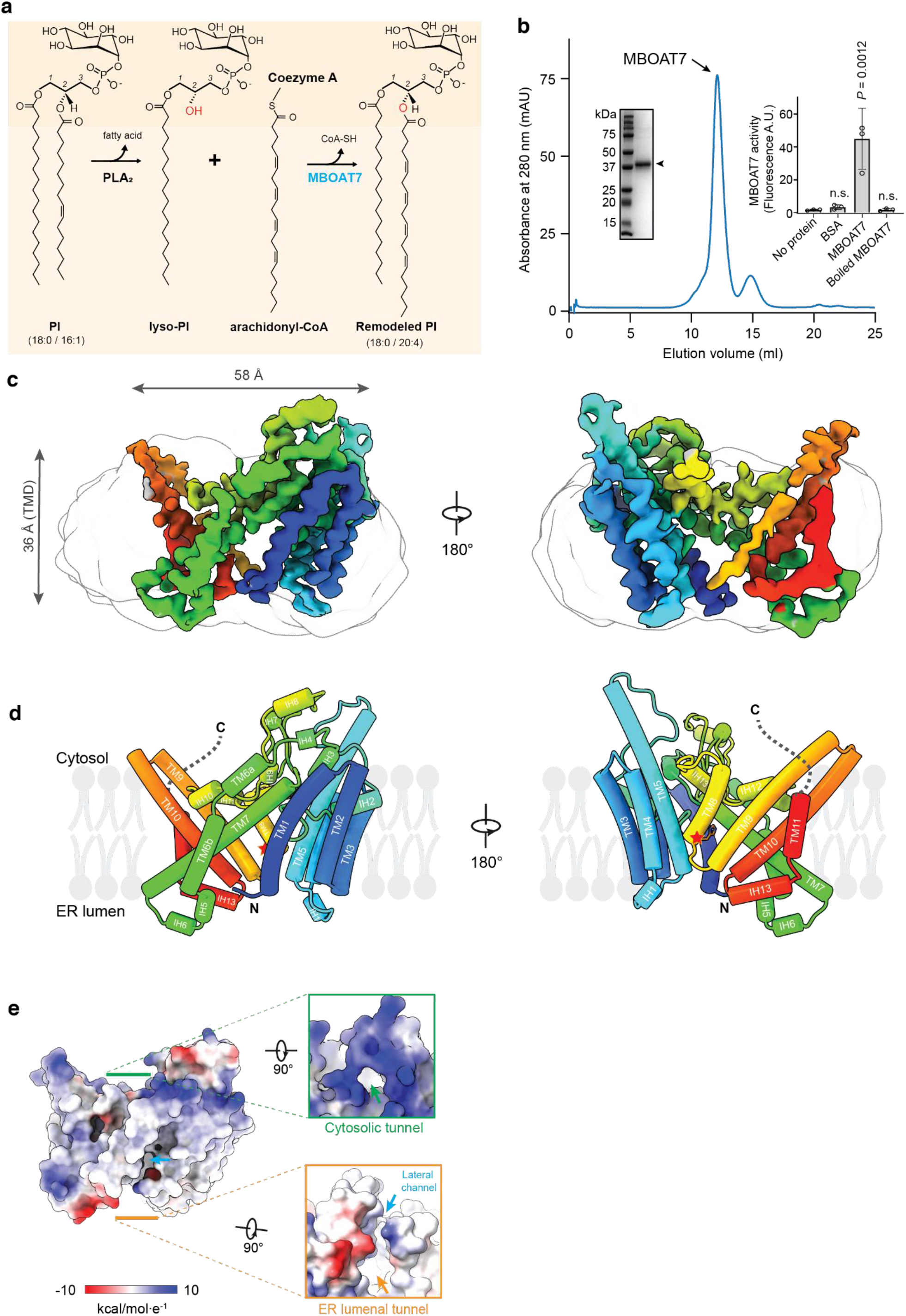
Cryo-EM structure of human MBOAT7. **a**, Diagram depicting the Lands cycle and MBOAT7-mediated PI remodeling in the membrane. Newly synthesized PI contains mainly monounsaturated *sn*-2 acyl chains. Phospholipase A2 (PLA2) hydrolyzes the *sn*-2 ester bond and produces lyso-PI. MBOAT7 specifically re-acylates lyso-PI with arachidonyl-CoA as the acyl donor to produce the remodeled PI molecules. The *sn*-2 hydroxyl group is highlighted in red. **b**, Size-exclusion profile of purified MBOAT7 reconstituted in PMAL-C8. Inset left, SDS-PAGE analysis of the MBOAT7 peak. The arrowhead denotes the MBOAT7 band, which migrates faster than its 53-kDa mass. Inset right, activity analysis of the purified MBOAT7 protein with the fluorescence monitoring of the production of the byproduct CoA-SH. **c,** Cryo-EM map of human MBOAT7. The amphipol micelle is shown as a transparent gray outline. Map is contoured at 6 σ. The map is rainbow-colored, corresponding the colors of secondary elements in the models in d. **d**, Cylinder representation of the human MBOAT7 model. The red star indicates the catalytic residue His356. Dashed lines indicate the disordered C-terminal region. TM, transmembrane; IH, intervening helix. **e**, Representation of MBOAT7 with an electrostatic surface. The inserts indicate the entries to the two tunnels from two sides of the membrane. Arrows indicate the specific features.

MBOATs form a large class of ER enzymes (11 in humans) that acylate small molecules, lipids, or proteins^8^. The structures of several MBOAT enzymes have been determined by cryo-electron microscopy (cryo-EM)^9–18^. Some MBOATs, such as porcupine and hedgehog acyltransferase, acylate polypeptides in the ER lumen^10,11,18^. Other MBOATs catalyze neutral lipid synthesis (e.g., DGAT1 and ACAT1/ACAT2) and act as multimers with a catalytic histidine residue that is buried deeply within the membrane^12–16^. For these MBOATs, the acyl-CoA substrate accesses the catalytic center through a tunnel that reaches from the cytoplasmic to the lumenal side of the ER. A lateral gate allows for entrance of the acyl acceptor substrate and exit of the neutral lipid product into the membrane.

There are fewer insights into the molecular mechanisms for MBOATs involved in the Lands cycle. The cryo-EM structure of chicken MBOAT5 (cMBOAT5) reveals a dimer, with each protomer having a fatty acyl–binding tunnel and a T-shaped chamber for esterification of arachidonyl-CoA with lyso-PC^9^. It remains unclear, however, how different MBOATs recognize different lysophospholipid substrates.

We focused our studies on MBOAT7 because of its unique function in PI metabolism and because understanding its catalytic mechanism has important implications for human physiology and disease. MBOAT7 deficiency in humans leads to developmental disorders of the central nervous system, with intellectual disability, epilepsy and autism spectrum disorder^19–21^. Deficiency of MBOAT7 function in liver is associated with increased lipogenesis in humans and is an important genetic risk factor for developing non-alcoholic fatty liver disease and other liver diseases^22–27^. Conversely, increased *MBOAT7* expression is correlated with detrimental outcomes in cancers, such as hepatocellular carcinoma and clear cell renal carcinomas^28,29^, suggesting that inhibitors of MBOAT7 may be useful therapeutic agents.

We determined MBOAT7’s molecular structure by single-particle cryo-EM. Combining structural insights with simulations and biochemical data, our findings led to a model for catalysis and substrate specificity of MBOATs. The MBOAT7 structure further enabled us to identify, by virtual screening, small-molecule lead compounds that specifically inhibit MBOAT7.

## Results

### Cryo-EM study of human MBOAT7 reveals characteristic structural features

To enable biochemical and structural analyses of human MBOAT7, we purified recombinantly expressed enzyme in detergent and reconstituted it into amphipol PMAL-C8 (Fig. 1b and Extended Data Fig. 1a). Purified MBOAT7 was similarly active in detergent or PMAL-C8 (Extended Data Fig. 1b). MBOAT7 in PMAL-C8 exhibited PI synthesis activity with a Vmax of ~300 pmol/min/μg protein, and Kms for arachidonyl-CoA and lyso-PI of 46 μM and 26 μM, respectively, which are similar to those of other MBOATs^10,12,30^ (Extended Data Fig. 1c,d). MBOAT7 was most active with arachidonyl-CoA (C20:4) as the acyl donor, followed by other unsaturated acyl-CoAs, and strongly preferred lyso-PI as the acyl acceptor (Extended Data Fig. 1e and f).

We used single-particle cryo-EM to obtain a density map of MBOAT7 at an overall resolution of ~3.7Å (Fig. 1c, Extended Data Fig.2), allowing us to build a model of the entire protein except the C-terminal region (442–472), which was not visible and presumably disordered. (Fig. 1d, Extended Data Fig.2). MBOAT7 comprises 11 transmembrane (TM) helical segments, connected by short intervening helices (IH) and loops, with the N- and C-termini on opposite sides of the membrane. Based on homology to other MBOATs, the N-terminus of MBOAT7 is predicted to localize to the lumenal side of the ER membrane. The monomeric protein occupies a space of ~36 × 58 × 28Å, embedded in the membrane (Fig. 1c) and is similar to the shape of a saddle.

The overall structure of MBOAT7 is similar to other MBOATs^9,10,12,14,17,18^. Two TM bundles (TM1–5, TM9–11) tilt away from each other, generating space in the center for the catalytic chamber. The chamber between these bundles is sealed by two long, diagonal TM segments (TM6 and TM7) on one side and TM8 on the other. The diagonal helix TM6 is broken into two short helices (TM6a and TM6b) (Fig. 1d); in contrast, a corresponding helix in cMBOAT5 is continuous^9^. TM8, containing the putative catalytic His356 residue, resides in the cavity between the two TM bundles. TM8 exhibit weaker density than the other TMs, indicating flexibility or multiple conformations in the apo state of the enzyme (Extended Data Fig. 2f).

Besides the “MBOAT core” (TM3–10, Extended Data Fig.3a), other regions of MBOAT7 are distinct from other MBOAT structures (Extended Data Fig.3b and c). For example, TM1-2 have features that are different from DGAT enzymes. MBOAT7’s N-terminal region contains only a short sequence before the first TM helix (TM1) and lacks sequences homologous to the longer N-terminal region of DGAT1, which mediates enzyme dimerization^12,13^. Also, DGAT1 enzymes have a large lateral gate that opens into the plane of the membrane, which presumably provides access for diacylglycerol substrates and an exit path for triacylglycerol products^12,13^. In contrast, TM helices 1 and 2 of MBOAT7 occlude the equivalent lateral gate region (Extended Data Fig.3d).

A tunnel twists through MBOAT7, connecting the cytoplasmic and the ER lumenal sides of the enzyme, with the active site located in the middle of this tunnel (Fig. 1e). This suggests a model similar to those proposed for other MBOATs^9–11^, in which acyl-CoA accesses the active site from the cytosolic side of the membrane and the acyl-acceptor, lyso-PI, enters the enzyme from the lumenal side. In MBOAT7, the lateral opening of the tunnel has an extended channel that branches off to the lumenal leaflet of the ER membrane.

### MBOAT7 model reveals how substrates access the enzyme active site

We were unable to obtain a high-resolution cryo-EM structure of MBOAT7 bound to substrates possibly due to increased sample heterogeneity. We, therefore, used the structure of cMBOAT5 bound to arachidonyl-CoA (PDB 7F3X) to model substrate binding^9^. These docking analyses revealed that arachidonyl-CoA fits well into the cytosolic part of the enzyme tunnel, positioning the thioester bond of the substrate near the catalytic His356 and Asn321 (Fig. 2a-c). Asn321 forms a hydrogen bond with the thioester carboxyl bond, implying it stabilizes the carbonyl group during the nucleophilic attack of His356 on the thioester bond. The adenine ring of the CoA moiety protrudes into the cytosol without strong interactions with the protein, similar to the acyl-CoA binding mode of other MBOAT enzymes^10,13,18^. The ribose group of CoA forms hydrogen bonds with Tyr333 (Fig. 2c), and the phosphate groups of CoA form salt bridges with Arg318. The pantetheine group has close contacts with Gln325, Met349 and Ser352, and mutation of these residues to alanine reduced the acylation activity of MBOAT7 (Fig. 2d).

**Fig. 2.**
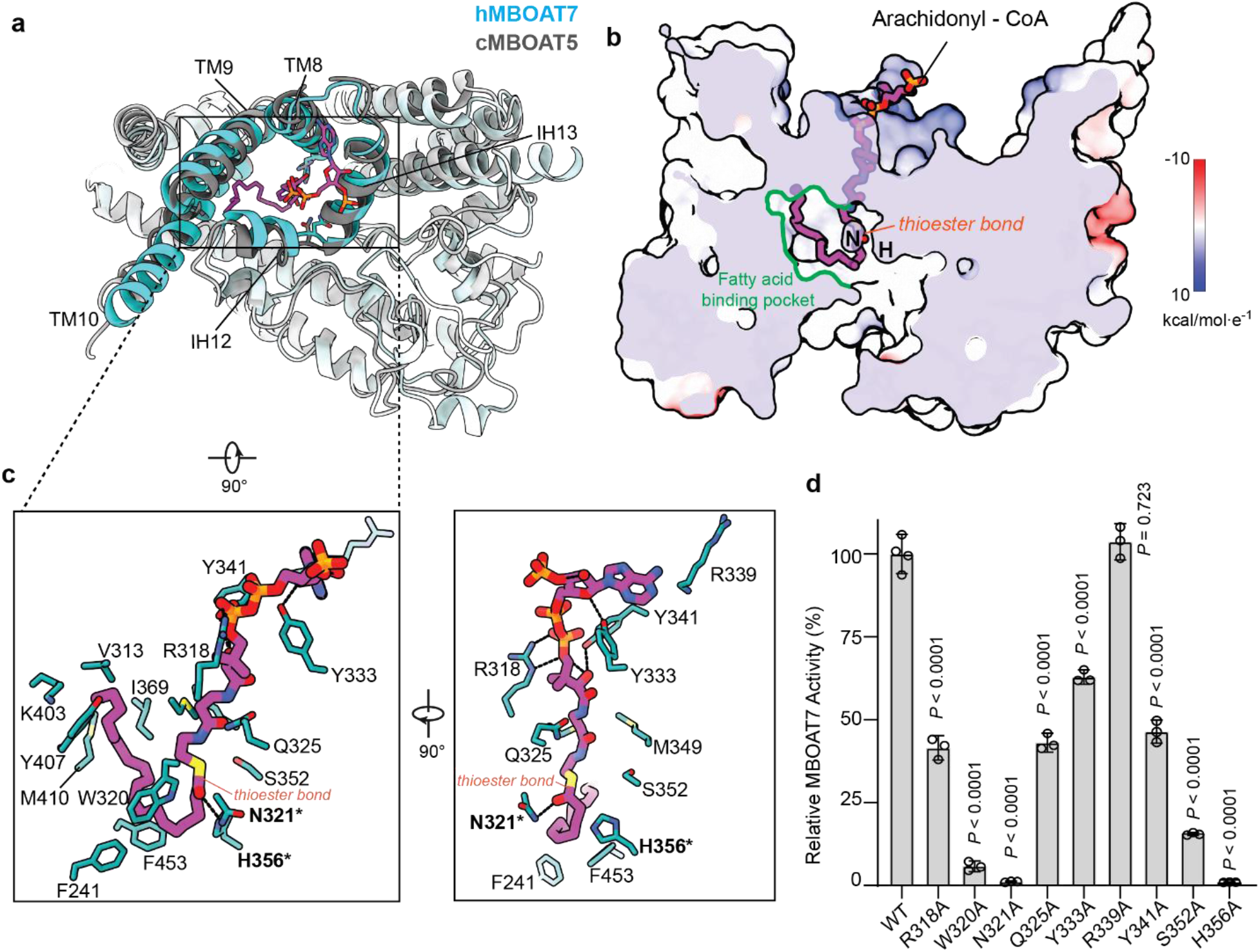
Arachidonyl-CoA accesses the MBOAT7 catalytic chamber via a tunnel from the cytosolic leaflet of the membrane. **a**, Top view of the arachidonyl-CoA molecule docked into the MBOAT7 cytosolic channel. Human MBOAT7 (hMBOAT7) structure is superimposed with the chicken MBOAT5 structure complexed with arachidonyl-CoA (PDB 7F40). The key helices making up the chamber, including TM8-10 and IH12-13, are highlighted in cyan. The root-mean-squared distance (RMSD) of 99 α-carbons within these regions between MBOAT5 and MBOAT7 is ~1.15Å. **b**, A cutting-in view of the catalytic chamber and side pocket bound with arachidonyl-CoA. MBOAT7 is represented with electrostatic surface. The positions of catalytic residues H356 and N321 are indicated by H and N, respectively. A twisted tunnel spans the enzyme from the cytosolic leaflet to the ER lumenal leaflet. The cytosolic access to the tunnel has a side pocket that accommodates arachidonyl-CoA binding. **c**, The interaction between arachidonyl-CoA and MBOAT7 residues. Polar interactions are shown with dashed lines. The catalytic residues are highlighted in bold with an asterisk. **d**, Activity of the arachidonyl-CoA binding channel alanine mutations. The enzymatic activities of mutants were normalized as the percentage of that of the wild-type MBOAT7 (mean ± SD, *n=3* independent experiments). Analysis was performed using one-way ANOVA with Dunnett’s *post hoc* test.

MBOAT7 strongly prefers arachidonyl-CoA as a substrate over other unsaturated fatty acyl CoAs (Extended Data Fig. 1e). The structure provides insight into this preference. The tunnel through MBOAT7 connects to a large cavity next to the catalytic residues, lined with bulky hydrophobic residues, including Phe241, Phe453 and Trp320 (the fatty acid–binding pocket in Fig. 2b, colored in green). Trp320 appears to form a hydrophobic gate to exclude water from the catalytic center, and mutating Trp320 to alanine strongly reduced MBOAT7 activity (Fig. 2b). The long, kinked acyl chain of arachidonyl-CoA fits well into this side cavity, interacting with Val313, Tyr407, and Ile369. In contrast, straight and saturated acyl-CoAs do not fit this bent cavity well, likely explaining the substrate preference of MBOAT7 for more flexible unsaturated acyl-CoAs.

MBOAT7’s tunnel provides a route for lyso-PI to enter the catalytic chamber from the lumenal side of the ER membrane. To gain insight into how specifically lyso-PI binds MBOAT7, we modeled lyso-PI (C18:0) binding to MBOAT7 and performed all atom-molecular dynamics (MD) simulations (Fig. 3). In the resultant energy-minimized structure, lyso-PI bound MBOAT7 within the ER lumenal opening of the tunnel. The inositol headgroup occupied the solvent-accessible part of the tunnel. The *sn-1* acyl chain extended into a hydrophobic side-channel formed by TM1, TM6b, and TM7 (Fig. 3a colored in cyan) that opens into the membrane. This channel is lined primarily with hydrophobic residues (e.g., Leu9, Phe205, Val247, Phe245) that align along the acyl chain of lyso-PI (Fig. 3d). In the apo-MBOAT7 structure, the width of the hydrophobic channel is narrow (~7Å in diameter) (Fig. 3b and c, hydrophobic channel is highlighted by cyan lines), suggesting that it cannot fit more than one acyl chain. This may explain how MBOAT7 preferentially binds lyso-PI with one acyl chain, but not the vasty more abundant PI with two acyl chains, which could result in product inhibition of the enzyme.

**Fig. 3.**
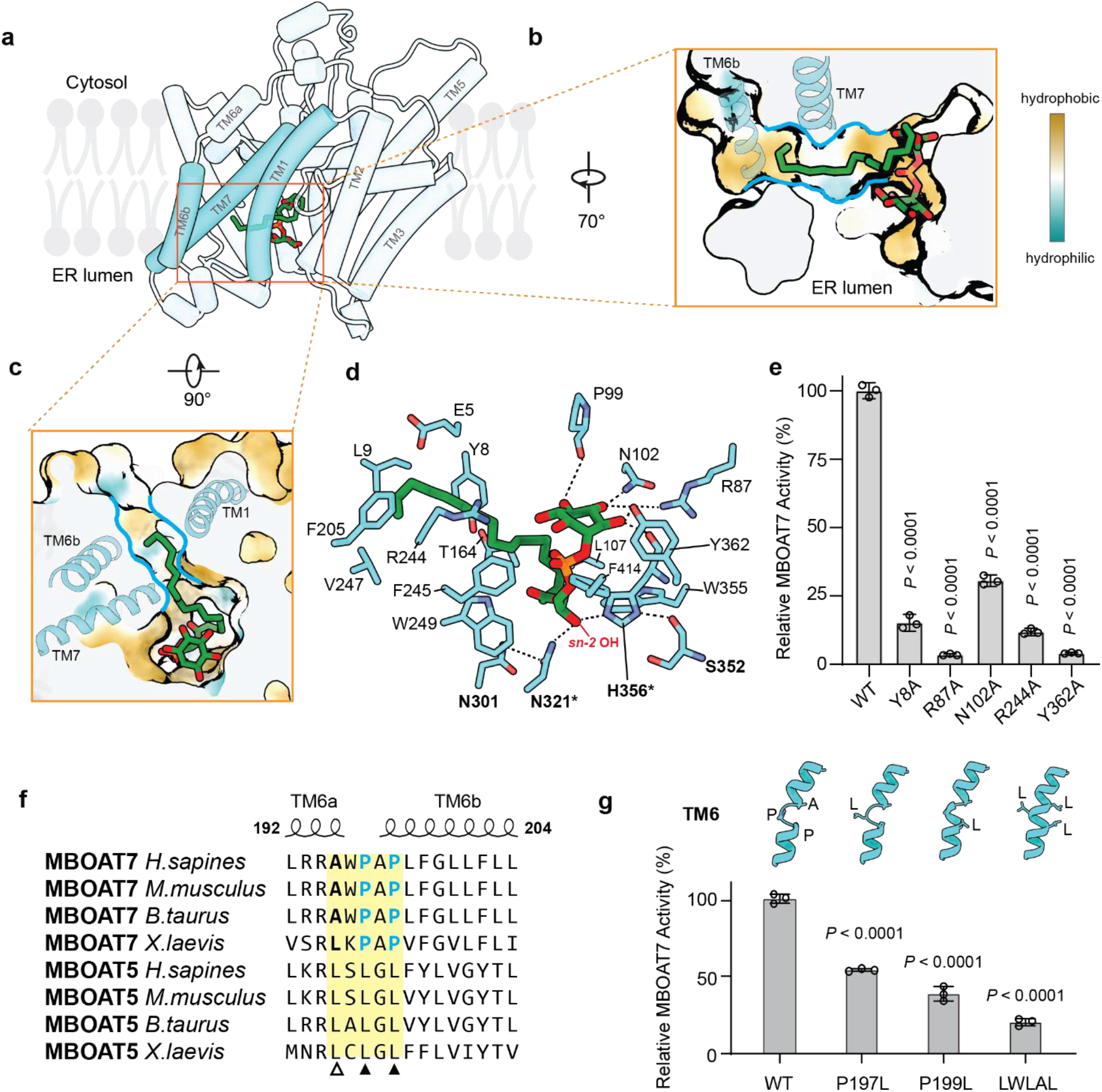
Lyso-PI accesses the MBOAT7 catalytic center through an ER lumenal opening. **a**, The lumenal access to the enzyme tunnel features an opening for the phospholipid headgroup and a hydrophobic channel for the acyl chain for lyso-PI binding. Cartoon representation of MBOAT7 with 18:0 lyso-PI at the energy minimal state of a molecular dynamic simulation. The helices making up the lateral channel are highlighted in cyan. **b** and **c**, Cutting-in views of the lyso-PI molecule in the catalytic chamber, viewed from the ER luminal side (b) or the membrane (c). MBOAT7 is represented with hydrophobicity surface. The TM helices comparing the lateral gate were shown in ribbons. **d**, Interaction of lyso-PI with MBOAT7 residues. The hydrogen bonds are shown with dashed lines. Residues involved in catalysis are shown in bold, and the two catalytic residues are indicated with asterisks. **e**, Activities of the inositol-binding residue alanine mutation. Enzymatic activities of mutants were normalized as the percentage of that of the wild-type MBOAT7 (mean ± SD, *n=3* independent experiments). Analysis was performed using one-way ANOVA with Dunnett’s *post hoc* test. **f**, Sequence alignment between MBOAT5 and MBOAT7 from different species. The secondary structure and residue numbers are shown from the human MBOAT7 apo structure model. The open triangle denotes the unconserved Ala195, while the solid triangles denote the conserved prolines unique to MBOAT7. **g**, Activity of the alanine mutations of the residues at the broken point of TM6. The enzymatic activities of mutants were normalized as the percentage of that of the wild-type MBOAT7 (mean ± SD, *n=3* independent experiments). Alphafold predictions on structures of MBOAT7 with the corresponding mutations. Ribbon representations of the TM6 are shown only.

In our structures, the catalytic residues His356 and Asn321 are close to the glycerol backbone (Fig. 3d). His356 can form hydrogen bonds with the hydroxyl group at the *sn*-2 position of the lyso-PI glycerol backbone and the backbone carbonyl group of Ser352 (Fig. 3d). The His356-Ser352 interaction likely increases the electronegativity of the imidazole Nε atom to facilitate the deprotonation of the *sn*-2 hydroxyl group of lyso-PI, as found in other MBOAT members^11,12^. Asn321 may also form a hydrogen bond with the *sn*-2 hydroxyl group of lyso-PI. Since it also hydrogen bonds with the arachidonyl-CoA thioester carbonyl group (Fig. 2c), Asn321 may assist in the acyltransferase reaction by stabilizing the two substrates close to catalytic His356.

### A discontinuous TM6 helix is important for MBOAT7 activity

Compared with other MBOAT structures, our cryo-EM structure of MBOAT7 is distinct, inasmuch as TM6 is divided into two short helices (Fig. 1d and Extended Data Fig. 3e) linked by two evolutionarily conserved proline residues that are not found in MBOAT1, 2 or 5 (Fig. 3f and Extended Data Fig. 6). Mutation of the MBOAT7 prolines resulted in a continuous helix in Alphafold2 predictions and markedly reduced lyso-PI acylation activity (Fig. 3g), suggesting that the discontinuous TM6 is required for optimal MBOAT7 activity.

We obtained a cryo-EM density map of MBOAT7 with lyso-PI and 18:1-ether CoA, a non-hydrolysable acyl-CoA analog (Extended Data Fig.7). This map provided adequate resolution (~ 6Å) to determine the overall architecture and TM arrangement of MBOAT7 with substrates. Docking this map with the apo-structure of MBOAT7 identified conformational changes in TM6: in the map obtained in the presence of substrates, TM6b moved away from the lateral portal, and TM6a moved down and became more horizontal. As a result, the lateral hydrophobic channel enlarged, as compared with the apo-structure (Extended Data Fig. 8). We hypothesize that this conformational change facilitates the release of the larger PI product with two acyl chains from the lateral portal, providing a possible explanation for how TM6 mutations reduced the enzyme activity.

### The region between TM4 and TM5 confers lysophospholipid substrate-specificity on MBOATs

The energy minimized structure suggests a unique recognition mode for the inositol headgroup. The inositol ring forms hydrogen bonds with two charged residues, Arg87 and Asn102, and the backbone carbonyl groups of Pro99 and His356. Two charged residues, Arg244 and Tyr8, are close to the phosphate and inositol groups and transiently interacted with these groups during MD simulations. Mutation of these residues decreased MBOAT7 catalytic activity (Fig. 3e). Besides Arg244, the phosphate group of lyso-PI is surrounded by a group of aromatic residues, including Tyr362, Phe414 and Phe245 and is stabilized by a combination of hydrogen bond and anion-pi interactions.

How MBOATs 1, 2, 5, and 7 specifically acylate lyso-PS, lyso-PE, lyso-PC, or lyso-PI is unclear. To investigate this, we docked three phospholipid headgroups as ligands into their corresponding MBOAT enzyme structures (obtained from this study or Alphafold predictions^31,32^). Each headgroup fits best into a pocket created by the converging ends of two highly variable TM bundles, containing parts of TM4, TM5, and the connecting loop (Fig. 4a and b), and a more conserved region spanning partial TM9 and TM10 (Fig. 4b). Important residues for recognizing the different headgroups are unique between different MBOATs and almost all clustered on the TM4 and TM5 variable regions (Fig. 4a–c). Specifically, the amino and carboxyl groups in the serine, representing the headgroup of lyso-PS, form salt bridges with three unique charged residues within the variable region in MBOAT1, including His122, Arg125 and Asp137. Choline, representing the headgroup of lyso-PC, is bound by Tyr129 and Cys146 in MBOAT5, a pattern that is similar to the interaction between cMBOAT5 and lyso-PC^9^. For MBOAT7, docking recapitulated Arg87 and Asn102 hydrogen bonding with the inositol headgroup (Fig. 3). Other important residues for headgroup recognition in TM9 and 10 are highly conserved and almost identical in the three MBOAT proteins analyzed (Fig. 4b and c).

**Fig. 4.**
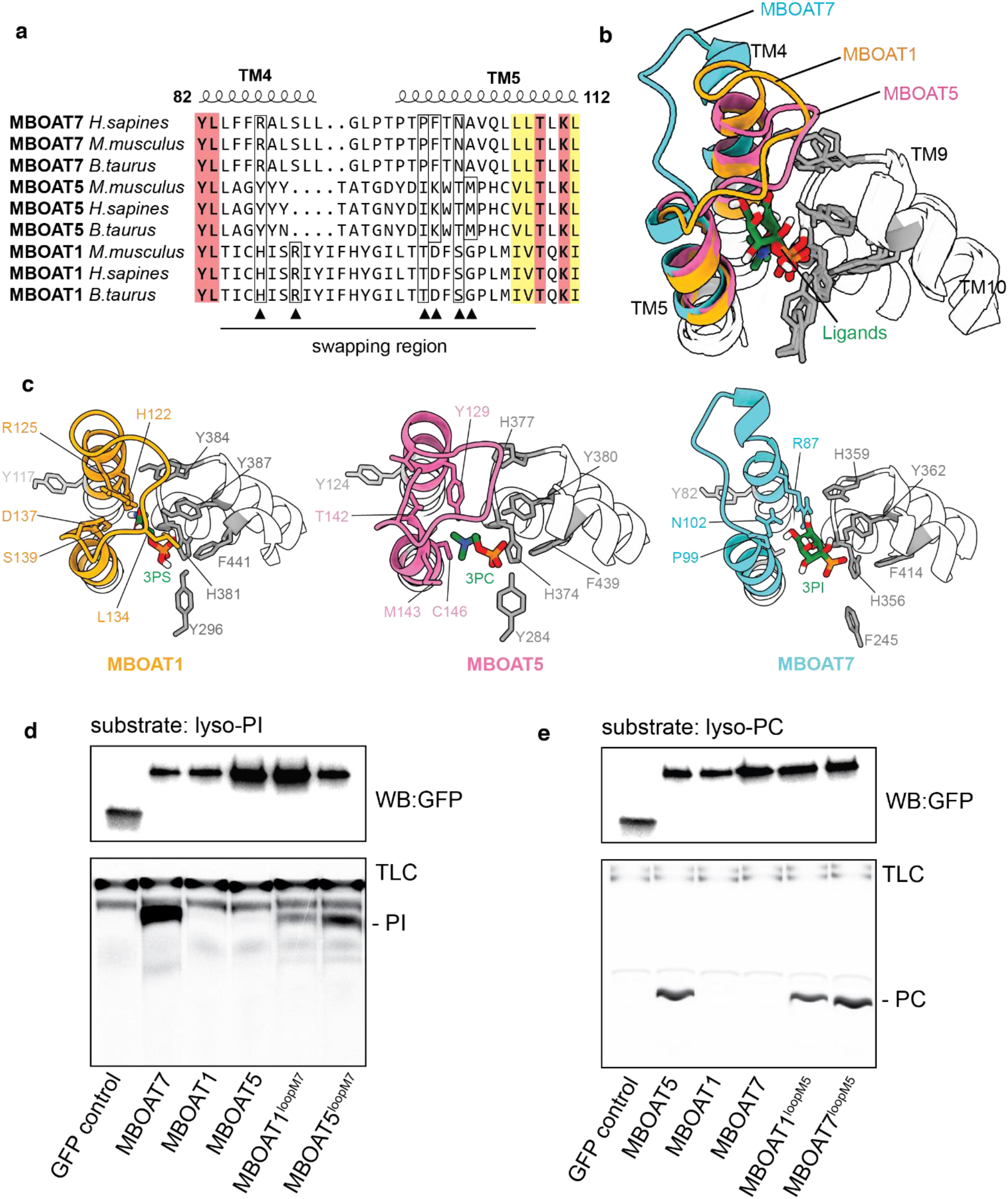
A variable region of MBOAT7 TM segments 4 and 5 determines the selectivity towards lysophospholipid substrates. **a**, Sequence alignment of MBOAT1, 5 and 7 from different species. Only the variable regions and the short flank sequences are shown. The full sequence alignment can be seen in Extended Data Fig. 6. Conserved residues are highlighted with colors and boldness. The key residues for recognizing the specific headgroups were denoted by frames and solid triangles. The swapping regions are also indicated. **b**, The overlay of pockets for the lyso-phospholipid headgroups from MBOAT1, 2 and 5. The highly conserved residues from TM9 and 10 are shown in grey, and the variable regions are shown in different colors. **c**, The detailed interaction between MBOAT and their specific phospholipid headgroups. The colors are consistent with b. 3PS, 3-phosphorylserine; 3PC, 3-phosphorylcholine; 3PI, 3-phosphorylinositol. **d** and **e**, The activity analysis of the region-swapped chimeric proteins. Protein levels were adjusted to the immunoblotting against GFP (upper). The activity is detected by the production of radioactive PI (d) or PC (e) on a TLC.

To test whether variable regions of TM4 and TM5 determine substrate specificity, we swapped these regions between MBOAT1, 5 and 7, and tested the activities of the chimeric MBOAT proteins (Fig. 4d and e). Upon replacing the variable regions of MBOAT1 and MBOAT5 with that of MBOAT7, these mutant enzymes gained acyltransferase activity towards lyso-PI (Fig. 4d). Similarly, replacing this region of MBOAT1 or MBOAT7 with that of MBOAT5 yielded enzymes that gained acyltransferase activity towards lyso-PC (Fig. 4d). Thus, these variable regions of MBOATs (corresponding to TM4 and TM5 of MBOAT7) constitute a specificity-determining region for different lyso-phospholipids.

### A structure-based screen identified MBOAT7 inhibitors

Increased *MBOAT7* expression correlates with detrimental outcomes in some cancers, including hepatocellular carcinoma and renal clear cell carcinomas, suggesting a dependency on the enzyme^28,29^. Genetic deletion of *MBOAT7* in renal clear carcinoma-derived cells induces cell-cycle arrest and prevents the cells from forming tumors *in vivo*^28^, suggesting MBOAT7-specific inhibitors may be valuable. To identify such inhibitors, we used VirtualFlow^33^, a structure-based virtual screening platform, to screen ~5.7 million commercially available compounds (Fig. 5a). We selected 12 candidate compounds with low free energy–docking scores and scaffold similarity and tested their effects on purified MBOAT7 activity at two concentrations, 50 and 5 μM (Fig. 5b). Two compounds, which we term Sevenin-1 and Sevenin-2, inhibited MBOAT7 activity. The IC50s for Sevenin-1 and Sevenin-2 were 0.87 μM [0.56 – 1.22 μM, 95% confidence interval (CI)] and 5.60 μM (4.61 – 6.82 μM, 95% CI), respectively (Fig. 5c). Notably, ATR-101^34^, a selective ACAT1 inhibitor, also inhibited MBOAT7 (Fig. 5b).

**Fig. 5.**
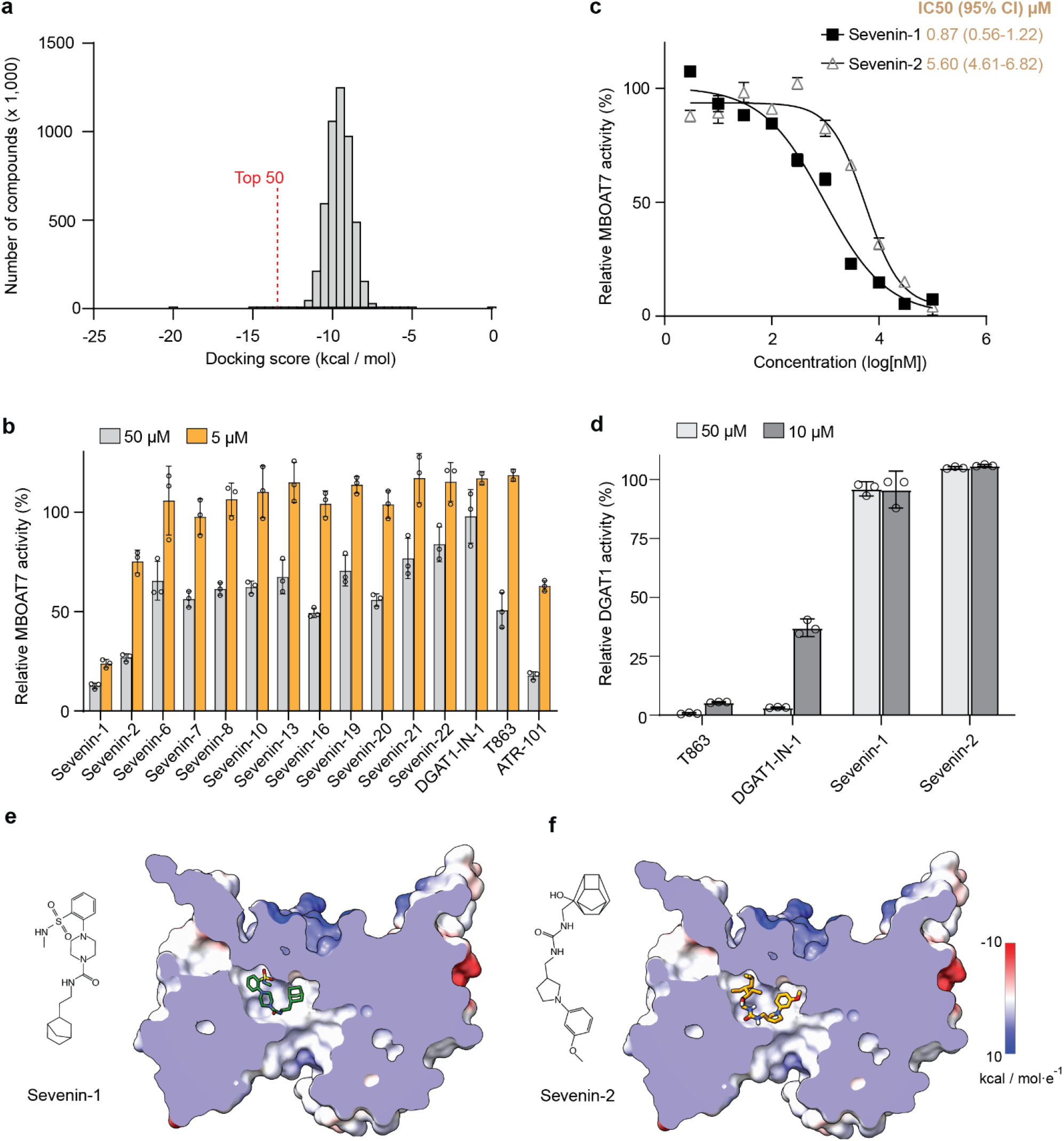
Structure-based discovery of MBOAT7 inhibitors. **a**, Histogram plot of the screen ~5.7 million compounds based on their free-energy docking score. **b**, Relative MBOAT7 activity with the addition of indicated compounds at 50 or 5 μM. Sevenin-number, compounds screened from VirturalFlow; the number indicates the compound ranking from the screen. Activities were normalized to vehicle control. **c**, IC50 curves for inhibition of MBOAT7 activity by Sevenin-1 and −2 measured by radioactivity-based assays (mean ± SD, *n=3* independent experiments). **d**, Inhibition of human DGAT1 activity by different compounds (mean ± SD, *n=3* independent experiments). **e** and **f**, Chemical structures of Sevenin-1 (**e**) and Sevenin-2 (**f**), and their docking positions in the MBOAT7 structure. MBOAT7 is shown with an electrostatic surface. Please note that the synthesized VF-2 is slightly different from the docked compound in VirtualFlow.

To screen for the specificity of Sevenin-1 and Sevenin-2, we measured their effects on purified human DGAT1^13^. Two DGAT1 inhibitors, T863 and DGAT1-IN-1, inhibited DGAT1 activity at 10 μM, whereas Sevenin-1 and −2 showed no inhibition even at 50 μM, suggesting that these two compounds may be MBOAT7-specific (Fig. 5d). According to docking results, both Sevenin-1 and −2 bind to MBOAT7 via occupying the acyl-CoA binding tunnel with the extended side pocket (Fig. 5e and f).

## Discussion

Here, we provide models for the structure of monomeric MBOAT7 and its catalytic mechanism to synthesize PI specifically from arachidonyl-CoA and lyso-PI substrates, and we identify chemical inhibitors of this reaction (Fig. 6). In this model, the arachidonyl-CoA substrate enters the enzyme tunnel from cytoplasmic leaflet of the ER. An extended lateral side pocket can accommodate the flexible acyl chain of unsaturated acyl-CoAs (Fig. 6a), but is not optimal for a straight, saturated acyl chains. Lyso-PI, enters the tunnel from the lumenal leaflet with a side channel harboring its single acyl chain and positions the *sn*-2 hydroxyl group of the glycerol backbone near the catalytic His356 residue.

**Fig. 6.**
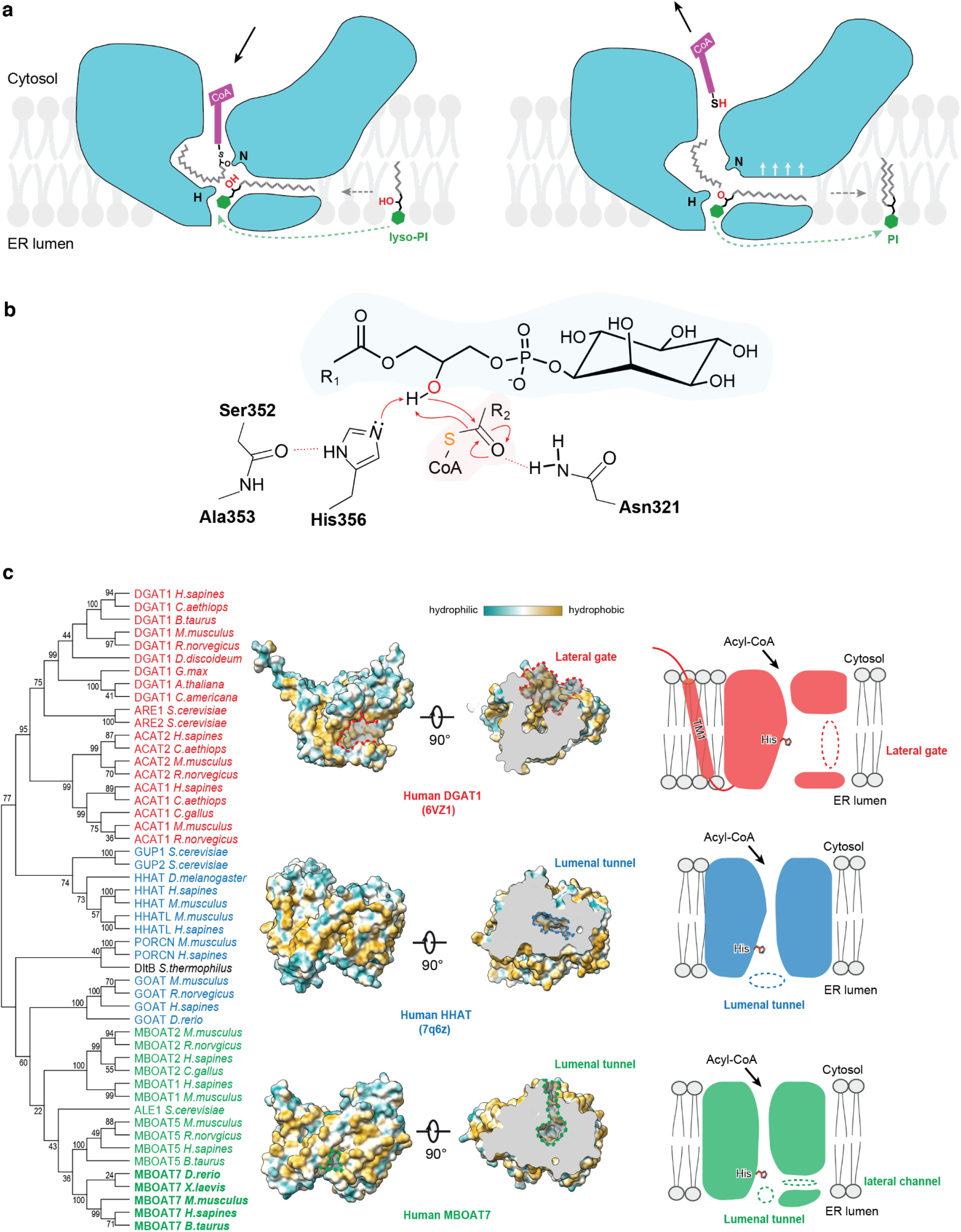
A model for MBOAT7 catalytic function and comparison of general MBOAT classes. **a**, Hypothetical model for MBOAT7-catalyzed PI remodeling. The membrane-partitioned or soluble acyl-donor arachidonyl-CoA enters the reaction center at the MBOAT7’s cytosolic tunnel. The side pocket accommodates the long and kinked arachidonyl acyl chain. The acyl-acceptor lyso-PI enters the catalytic chamber through the connected lateral channel and ER lumenal tunnel, with the hydrophobic acyl chains enters through the lateral channel (grey dashed arrow), and the hydrophilic inositol headgroup slides in through the ER luminal tunnel (green dashed arrow). The thioester bond and *sn*-2 hydroxy group of lyso-PI are positioned close to the catalytic Asn321 (N) and His356 (H). Once the catalysis is complete, CoA-SH is released into the cytosol, and PI is released to the ER luminal leaflet of the membrane. The lateral gate is opened slightly wider to facilitate the rapid release of PI. **b**, Putative chemical mechanism of MBOAT7-catalyzed PI remodeling. The backbone carbonyl of Ser352 forms hydrogen bonds with the N on the imidazole ring, which polarizes the other N to deprotonate the sn-2 hydroxyl group of lyso-PI glycerol. Deprotonated hydroxyl initiates a nucleophilic attack on the thioester bond of the arachidonyl-CoA. Finally, CoA-S^−^ attracts the proton back from His356 to complete the catalytic cycle. **c**, Phylogenetic analysis and structural comparison of MBOATs in different clusters. MBOATs with different substrate preferences were clustered and highlighted with different colors except the bacteria DltB. Red, blue and green indicate preference towards neutral lipids, polypeptide, and lyso-phospholipids, respectively. One representative structure from each cluster is shown with hydrophobicity surfaces. The key features for acyl acceptors to access are highlighted in dashed lines and labels. Cartoons are also shown on the right. Dashed circles denote the different entry sites of the acyl-acceptor substrates.

In the putative catalytic mechanism, His356 deprotonates the hydroxyl group of the glycerol backbone, enabling the oxygen to undergo a nucleophilic attack on the carbonyl group of the thioester bond (Fig. 6b). After forming an intermediate, electron transfers would result in formation of the CoA-SH and PI products. We hypothesize that binding of the substrates causes conformational changes in the TM6 helix to widen the hydrophobic, lumenal opening of the enzyme, allowing release of PI, with two acyl chains, into the membrane.

A key question for MBOAT enzymology is how substrate specificity is achieved. Our data suggest that TM segments 4–5 of MBOAT7 (or equivalent sequences of other Lands cycle MBOATs) constitute a lysophospholipid “specificity-determining region”. Key residues in this region are conserved for each different MBOAT and appear to interact with each headgroup (Fig. 4).

Beyond Lands cycle enzymes, phylogenetic analyses of MBOATs clusters them into groups according to their substrates (e.g., hydrophobic neutral lipids, hydrophilic polypeptides, or amphipathic lyso-phospholipids) (Fig. 6c). Whereas the acyl-CoA binding regions of MBOATs are relatively conserved, different paths appear to allow acyl-acceptor substrates to access the catalytic site (Fig. 6c). For MBOATs that acylate proteins (e.g., hedgehog acyltransferase, porcupine), the protein substrate reaches the catalytic center from the ER lumen. For MBOATs that catalyze neutral lipid synthesis (e.g., DGAT1, ACAT1, ACAT2), the substrate accesses the catalytic center from a lateral gate in the plane of the membrane. For the MBOATs that mediate phospholipid remodeling, the lysophospholipid substrates access the enzyme tunnel from the lumenal side.

Loss-of-function mutations in human *MBOAT7* result in liver disease, intellectual disability, early onset seizures, and autism spectrum disorders^19–21,24–27,29^. Mapping the pathogenic *MBOAT7* mutations responsible for these disorders on to our MBOAT7 structure model enables some predictions on how they disrupt MBOAT7 function (Extended Data Fig. 9). For example, the E253K disease-causing mutation^19^, though located in TM7 away from the catalytic center, may disrupt hydrogen bonds with a nearby loop and destabilize the protein fold. The R384Q mutation may interfere with binding of acyl-CoA substrates.

The MBOAT7 structure enabled identification of two potential inhibitors, Sevenin-1 and - 2 (Fig. 5). Although these compounds require further characterization and optimization, they provide a foundation for developing selective MBOAT7 inhibitors as therapeutic agents for diseases with increased MBOAT7 activity, such as cancers, with MBOAT7 dependency. Interestingly, similar to other MBOAT inhibitors^18,35,36^, the structures of Sevenin-1 and −2 contain two bulky aromatic groups connected by an amide bond. This could be a common scaffold for the MBOAT inhibitors, which may help guide the design of future MBOAT inhibitors.

## Acknowledgments

We thank S. Sterling and M. Mayer at the Harvard cryo-EM center, and C. Xu and K. Song at the University of Massachusetts cryo-EM facility, for electron microscopy data collections. The MD simulations were performed on the high-performance GPU cluster (GM4) at the University of Chicago Research Computing Center (supported by NSF grant DMR-182869). The VirtualFlow drug screen was performed using the high-performance CPU clusters at the Harvard Medical School O2 system. We are grateful for discussions with the members of the Liao and Farese & Walther laboratories, and we thank Gary Howard for editorial assistance. S.K. and G.A.V. were supported by R01GM063796. This work was supported by R01GM141050 (to R.V.F) and the Howard Hughes Medical Institute (T.C.W).

## Author Contributions

K.W., M.L., T.C.W. and R.V.F. conceived the project. K.W. performed protein expression, purification, cryo-EM grid preparation, data processing, model building, molecular docking and VirtualFlow drug screen studies. M.L. and S.W. advised on cryo-EM data processing. K.W., C.L., and A.B.H. performed the mutagenesis and activity studies. S.K. and G.A.V. performed molecular dynamics simulations. A.J.B. performed the mass spectrometry analysis. K.W., M.L., T.C.W. and R.V.F. wrote the manuscript. All authors analyzed and discussed the results and contributed to the manuscript.

## Methods

### Protein expression and purification

MBOAT7 from *Homo sapiens* (UniPort ID: Q96N96) was expressed in HEK293 freestyle cells, using the inducible stable cell line system^37^. Briefly, the MBOAT7 cDNA sequence was cloned into the pSBtet vector with a C-terminal GFP-strepII sequence. The stable cell line was generated by co-transfection of the pSBtet-MBOAT7 and the SB-100X transpose vector, followed by a week of puromycin selection. The stable cells were grown in suspension at 37 °C in FreeStyle 293 Expression Medium (ThermoFisher). When the cells reached a density of 2.0–2.5 × 106 cells per ml, 20 μM doxycycline and 5 mM sodium butyrate were added to the cells to induce the protein production and boost the expression, respectively. Temperature was set to 30 °C. The cultures were harvested after ~60 hours. The cells were washed once in ice-cold PBS, then flash frozen in liquid nitrogen and stored in −80 °C or placed on ice for immediate use.

All protein purification steps were performed at 4 °C. Thawed cell pellets were re-suspended in the lysis buffer containing 50 mM Tris-HCL, pH 8.0, 150 mM NaCl, 1 mM EDTA and supplemented with 1 × complete protease inhibitor cocktail (Roche). DDM/CHS detergent (1%/0.1%) (Anatrace) was added to lyse the cells for 2 hours. Insoluble debris was removed by centrifugation 50,000 × *g* for 45 minutes, and the supernatant was incubated with prewashed Strep-Tactin Sepharose beads (IBA) for 1 hour. The resin was then washed with ~20 ml of washing buffer (lysis buffer containing 0.1% digitonin) before the elution of GFP-fused MBOAT7 protein by washing buffer containing 2 g/L desthiobiotin. TEV proteases were then added to the eluted proteins to remove the GFP and strep tag before the elution was concentrated by a 50-kDa MWCO spin concentrator and injected onto a Superose 6 column (GE Healthcare) equilibrated in 100 mM Tris-HCl, pH 7.5, 150 mM NaCl, and 0.05% (w/v) digitonin. The peak fractions containing MBOAT7 were pooled and concentrated. To reconstitute MBOAT7 proteins in PMAL-C8 (Anatrace), MBOAT7 in detergent was mixed with PMAL-C8 at a 1:3 (w:w) ratio, followed by gentle agitation overnight in the cold room. Detergent was then removed by mixing with Bio-Beads SM-2 (Bio-Rad) for 1 hour. Bio-beads were removed by passing through a disposable polyprep column. The MBOAT7 protein was further purified by injecting onto a Superdex 200 increase 10/300 GL column equilibrated with 100 mM Tris-HCl, pH 7.5, and 150 mM NaCl.

To prepare the protein sample with lyso-PI and acyl-CoA, 10 μM lyso-PI was added at each step after cell solubilization and before PMAL-C8 reconstitution, whereas 10 μM acyl-CoA or nonhydrolyzable analogs was added at each step before the final size-exclusion purification.

### Cryo-EM imaging and data processing

Grids for cryo-EM imaging were prepared on a Vitrobot Mark IV system (ThermoFisher Scientific) by applying 3 μL of PMAL reconstituted MBOAT7 at 5–6 mg/ml to Quantifoil holey gold grids (Au R1.2/1.3; 400 mesh) that were glow discharged for 30 seconds at 15 mA right before use. Grids were blotted for 5 seconds at 4 °C, 100 % humidity, and a blot-force of 12 N before being plunge frozen in liquid ethane cooled by liquid nitrogen. Cryo-EM data were collected on a Titan Krios electron microscope with a K3 direct electron detector. Movies were collected in counting mode with a pixel size of 0.825Å and a total does 50.4 electron/Å2.

Movies were corrected for beam-induced motion by MotionCor2^38^, and contrast transfer function parameters were determined using CTFFIND4^39^. Template particle picking and 2D classification were performed using Simplified Application Managing Utilities of EM Labs (SAMUEL) similarly to previous dataset processing^40^. To identify particles with rare angles, the deep-learning-based particle picker Topaz^41^ was used and followed by multiple rounds of 2D classifications in cryoSPARC^42^. Selected particles from SAMUEL and cryoSPARC were merged, and the redundant particles were removed before 3D processing with Relion3.0^43^, including 3D classification, 3D refinement, and domain masking. 3D classification on binned particles at 3.3Å pixel size were first performed to remove bad particles, and additional rounds of 3D classification were performed on less binned and unbinned particles. 3D classes were combined or discarded based on the quality of cryo-EM density of the transmembrane helices. Overall resolution of cryo-EM map was computed according to the gold standard Fourier shell correlation method. The local resolution was calculated with the local resolution estimation function in Phenix^44^. A flow-chart summary on the data processing can be found in Extended data Figure 2. The statistical details related to data processing are summarized in Supplemental Table 1.

### Model building and refinement

The Alphafold 2 predicted MBOAT7 model was used as the initial model. The model was docked into the map and refined using phenix.real_space_refine in Phenix^44^ and manually adjusted in COOT^45^. The resulting map was further put back through the real-space refinement procedure to undergo further refinement. The final model was validated with MolProbility^46^.

### Mass spectrometry

To confirm substrate binding with purified MBOAT7 during protein preparation, lipid contents were extracted using the tert-butyl-methyl ether and methanol method^47^ from purified MBOAT7 proteins and lipid controls (Avanti Polar Lipids). Extracted lipids were resolubilized in 300 μL of isopropanol, methanol and chloroform (4:2:1, v:v:v) containing 7.5 mM ammonium acetate. Mass spectrometry analysis of lipid mixture was performed using an Orbitrap ID-X Tribrid mass spectrometer (ThermoFisher Scientific), equipped with an automated Triversa Nanomate nanospray interface (Advion Bioscience) for direct infusion lipid solution delivery. All full scan mass spectra were acquired in negative mode using mass resolution of 500,000 (FWHM at m/z 200). Visualization of mass spectrometry data was performed using Qual Browser on Xcalibur 4.3.73.11 software (Thermo Scientific).

### Molecular dynamics simulation and docking

MBOAT7 was placed in a POPC bilayer. The topology of lyso-PI was made based on the topology of PI and diacylglycerol and is available at git@github.com:ksy141/TG.git. Simulations were carried out three times, each of which was 500 ns long. Simulations were performed using the GROMACS (version 2018)^48^ simulation engine with the CHARMM36m lipid and protein force field^49,50^. Simulations were integrated with a 2-fs timestep. Lennard-Jones pair interactions were cut off at 12Å with a force-switching function between 10 and 12Å. The Particle Mesh Ewald algorithm was used to evaluate long-range electrostatic interactions.^51^ The LINCS algorithm was used to constrain every bond involving a hydrogen atom.^52^ A temperature of 310 K and a pressure of 1 bar were maintained with the Nose-Hoover thermostat and the Parrinello-Rahman barostat, respectively.^53–55^ The coupling time constants of 1 and 5 ps were used, respectively. A compressibility of 4.5 × 10-5 bar-1 was used for semi-isotropic pressure coupling. MD Analysis was used for analyzing the trajectories^56^.

The constraint docking was performed using the MedusaDock 2.0 program^57^. The headgroup search space were estimated based on cMBOAT5 and MBOAT7 structures. Constraints were set between the phosphate group and conserved interacting residues.

### MBOAT7 mutagenesis and activity assays

The Michaelis-Menten and substrate specificity assays were performed using purified and PMAL reconstituted MBOAT7 with a fluorescence-based coupled-enzyme assay, as described^12,14^. Essentially, the activity was monitored by detecting the release of the free CoA-SH with the fluorescence probe 7-diethylamino-3-(4-maleimidophenyl)-4-methylcoumarin (CPM). A 10-μL reaction mixture contains 2 mg/ml BSA, 75 mM Tris, pH 7.5, 100 nM methyl arachidonyl fluorophosphonate (MAFP), 150 mM NaCl, and 0.025 g/L of MBOAT7 protein and the substrates acyl-CoA and lyso-PI with varied concentrations. For Michaelis-Menten characterization, either arachidonyl-CoA or lyso-PI was maintained at 100 μM, and the concentrations other substrate were varied. For characterizing substrate specificity, different acyl-CoA and lyso-phospholipids were added to a final concentration of 100 μM. The reaction was initiated by adding the enzymes to the reaction. The reaction proceeded for 3 minutes at 37 °C before being stopped by adding 1.3% SDS and incubating at room temperature for 5 minutes. 90 μL of CPM working solution (50 μM in 75 mM Tris, pH 7.5 and 150 mM NaCl) was added to the reaction, and the mixture was kept at room temperature for 30 minutes and followed by fluorescence detection (excitation 355 nm; emission 460) using the Tecan i-control infinite 200 plate reader. The background was determined from reactions without adding enzymes and subtracted to remove the background noise. Nonlinear regression to the Michaelis-Menten equation and allosteric sigmoidal analysis were performed using GraphPad Prism 9.

MBOAT7 mutants were generated by site-directed mutagenesis on the pFBM construct expressing MBOAT7 with a C-terminal GFP and strep tag using the Q5 Mutagenesis Kit (New England Biolabs), according to the manufacturer’s protocol. The sequenced constructs were transfected into the HEK293 freestyle cells with PEI, and the cells were harvested after 70 hours incubation at 30 °C. The cells were lysed and proteins were purified as described above with the modification that DDM/CHS was used for all solubilization, purification, and elution steps, and 10% glycerol was added in the final elution buffer. The initial levels of the wild-type and mutant MBOAT7 proteins were measured by immunoblotting against the GFP tag fused to the C-terminal of MBOAT7 and adjusted to the same quantity for assays. The acyltransferase activity was determined by measuring the incorporation of radiolabeled oleoyl-CoA into the product phosphatidylinositol^6^. In brief, a total 100 μL reaction mixture was prepared containing 75 mM Tris-HCl, pH 7.4, 2 g/L delipidated BSA, 150 mM NaCl, 100 nM MAFP, 50 μM 18:0 lyso-PI, 100 μM oleoyl-CoA containing 0.2 μCi [^14^C]-oleoyl-CoA as a tracer (American Radiolabeled Chemicals), and proteins (free GFP control, WT or mutant MBOAT7, ~ 100 ng) to initiate the reaction. The reaction was started by adding proteins to the reaction mixture and incubated for 15 minutes at 37 °C. The reactions were quenched by adding chloroform/methanol (2:1 v:v), followed by 2% phosphoric acid for phase separation. The organic phase was harvested, dried, resuspended in chloroform and loaded on a silica gel TLC plate (Analtech). Lipids were resolved in a solvent system consisting of chloroform, methanol, acetic acid, and water (65:35:8:5 v:v:v:v). The radioactivity of the phosphatidylinositol bands was revealed by Typhoon FLA 7000 phosphor imager (GE Healthcare Life Sciences) and quantified by Quantity One (V4.6.6).

### VirtualFlow drug screening

The structure-based drug screen was performed essentially according to the instructions from VirtualFlow website (https://virtual-flow.org/) and github page (https://github.com/VirtualFlow/VFVS). The searching space was determined by covering both the cytosolic and ER lumenal tunnels of MBOAT7 using the AutoDockTools^58^. 5.7 million ligands were selected from the REAL library with molecular masses 375–425 Daltons. The VirturalFlow screen was performed using QuickVina program, and the final 50 compounds were further validated using the Autodock program^58^. Twelve compounds were further demanded from the Enamine company and tested *in vitro*. For some compounds, the chemical structures of the synthesized compounds and original docked compounds in VirtualFlow are slightly different possibly due to the file conversion during library preparation or library update in Enamine.

### Data availability

The three-dimensional cryo-EM density maps have been deposited into the Electron Microscopy Data Bank under accession numbers EMD-XXX (apo) and EMD-XXX (with substrates). The coordinates are deposited into the Protein Data Bank with accession numbers XXXX (apo). All other un-deposited model and data are available upon request.

---

**Extended Data Table 1.** Cryo-EM data collection, refinement and validation statistics.

|  | MBOAT7 in PMAL-C8<br>(EMDB-xxxx)<br>(PDB xxxx) | MBOAT7 in PMAL-C8 with lyso-PI<br>and 18:0 ether CoA<br>(EMDB-xxxx) |
| --- | --- | --- |
| **Data collection and processing** |  |  |
| Magnification | 105,000 | 105,000 |
| Voltage (kV) | 300 | 300 |
| Electron exposure (e-/Å²) | 50 | 50 |
| Defocus range (µm) | 0.8 – 2.2 | 1.1 – 2.5 |
| Pixel size (Å) | 0.825 | 0.825 |
| Symmetry imposed | C1 | C1 |
| Initial particle images (no.) | 25,824,808 | 3,730,800 |
| Final particle images (no.) | 206,418 | 289,236 |
| Map resolution (Å) | 3.7 | 6.0 |
| FSC threshold | 0.143 | 0.143 |
| Map resolution range (Å) | 2.8 – 10.9 | 4.9 – 15.3 |
| **Refinement** |  |  |
| Initial model used | AF-Q96N66-F1-model_v2<br>(AlphaFold2 model) |  |
| Model resolution (Å) | 3.18 |  |
| FSC threshold | 0.143 |  |
| Model resolution range (Å) | 211.2 - 3.0 |  |
| Map sharpening *B* factor (Å²) | -120 |  |
| Model composition |  |  |
| Non-hydrogen atoms | 3510 |  |
| Protein residues | 441 |  |
| *B* factors (Å²) |  |  |
| Protein | 105 |  |
| R.m.s. deviations |  |  |
| Bond lengths (Å) | 0.003 |  |
| Bond angles (°) | 0.671 |  |
| Validation |  |  |
| MolProbity score | 2.1 |  |
| Clashscore | 8.11 |  |
| Poor rotamers (%) | 0 |  |
| Ramachandran plot |  |  |
| Favored (%) | 97.72 |  |
| Allowed (%) | 2.28 |  |
| Disallowed (%) | 0 |  |

**Extended Data Fig. 1.**
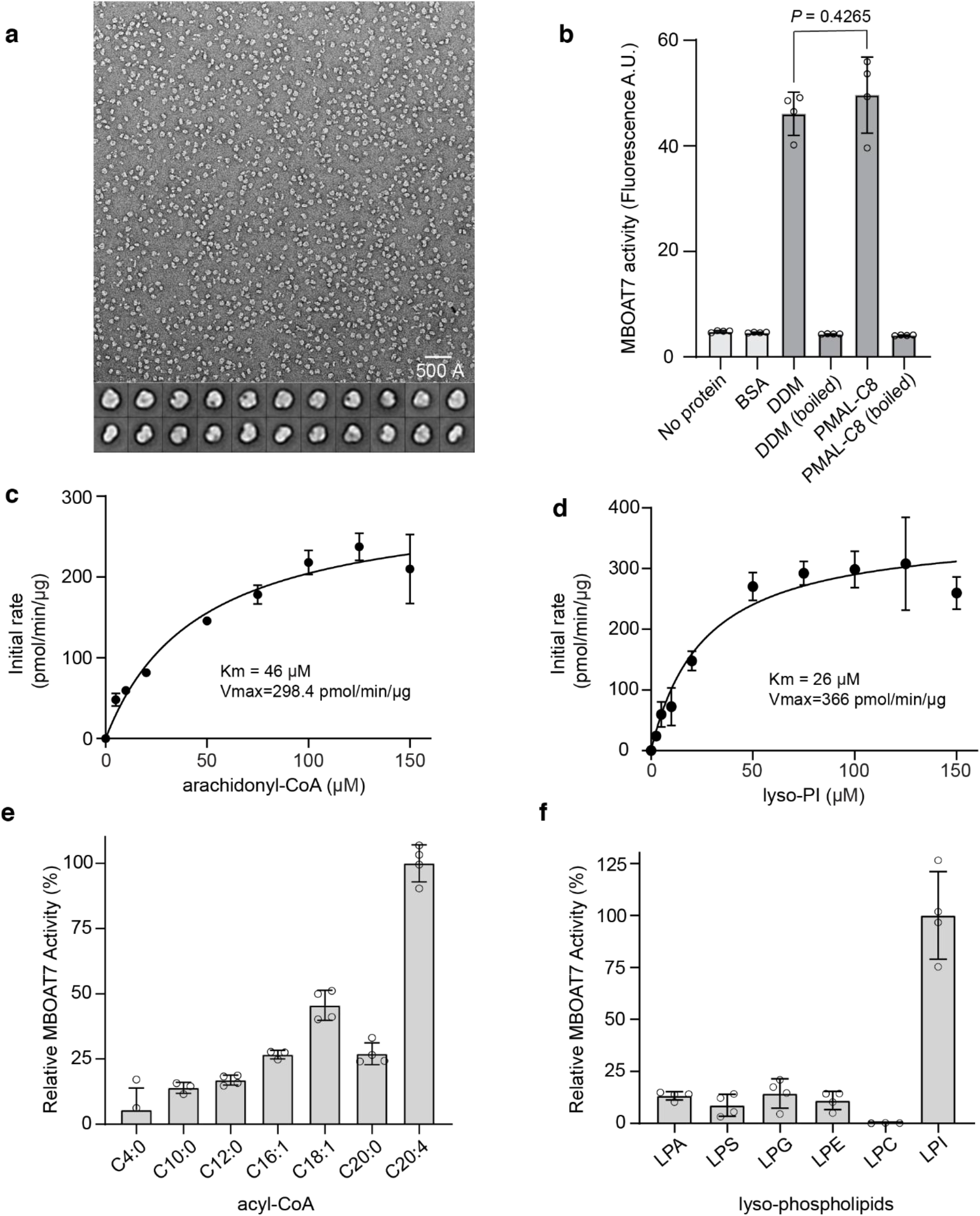
Characterization of purified human MBOAT7. **a,** Representative negative-stain electron micrograph and 2D averages of purified MBOAT7 in PMAL-C8. **b**, Activity comparison of purified MBOAT7 in DDM detergent or PMAL-C8. The activity is quantified by monitoring the fluorescent production of Co-SH that is covalently linked to CPM (mean ± SD, *n*=4 independent experiments). Analysis was performed using unpaired *t* test. **c** and **d**, Initial rate of reaction versus two substrates, arachidonyl-CoA (c) and lyso-PI (d). *n*=3 independent experiments. **e**, MBOAT7 activity towards acyl-CoA with different acyl chains (number of carbons: number of double bonds). *n*=4 independent experiments. **f,** MBOAT7 activity towards different lyso-phospholipids. *n*=4 independent experiments. LPA, lyso-phosphatidic acid; LPS, lyso-phosphatidylserine; LPG, lyso-phosphatidylglycerol; LPE, lyso-phosphatidylethanolamine; LPC, lyso-phosphatidylcholine; LPI, lyso-phosphatidylinositol.

**Extended Data Fig. 2.**
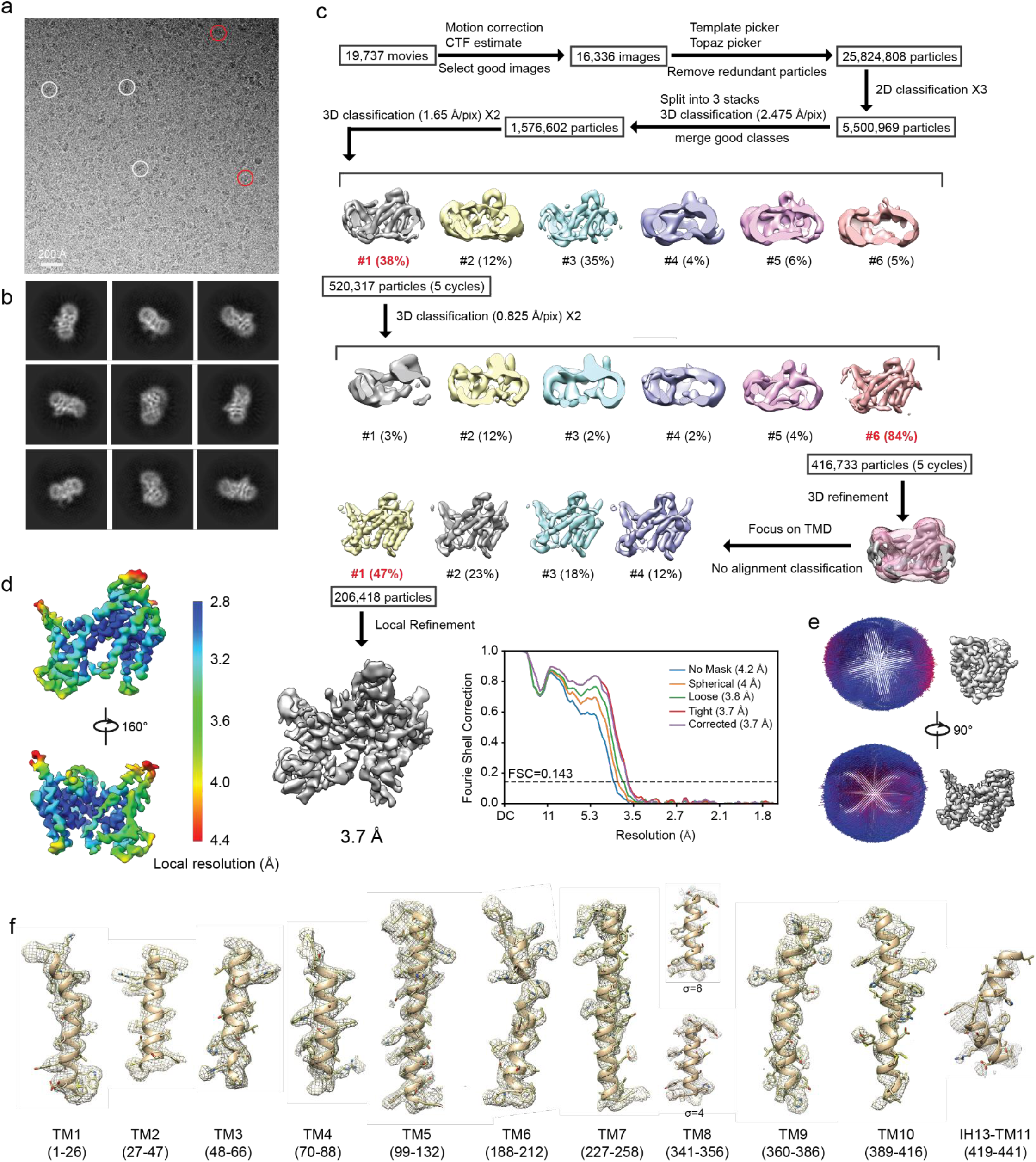
Cryo-EM image processing of human MBOAT7 in PMAL-C8. **a,** Representative cryo-EM image of MBOAT7. Some particles are highlighted with white (side views) or red (end-on views) circles. **b**, Representative 2D class averages. **c**, A flow chart for data processing (see Methods for details) and its FSC curves between two half maps with indicated resolutions at FSC=0.143. **d**, Local-resolution map of MBOAT7 in two orientations. **e**, Angular distribution of particle images included in the final 3D reconstruction. f, Cryo-EM densities superimposed with atomic model for individual transmembrane helices (TM1-TM11). Maps are contoured at 6 σ, except TM8 which is shown at two contour levels.

**Extended Data Fig. 3.**
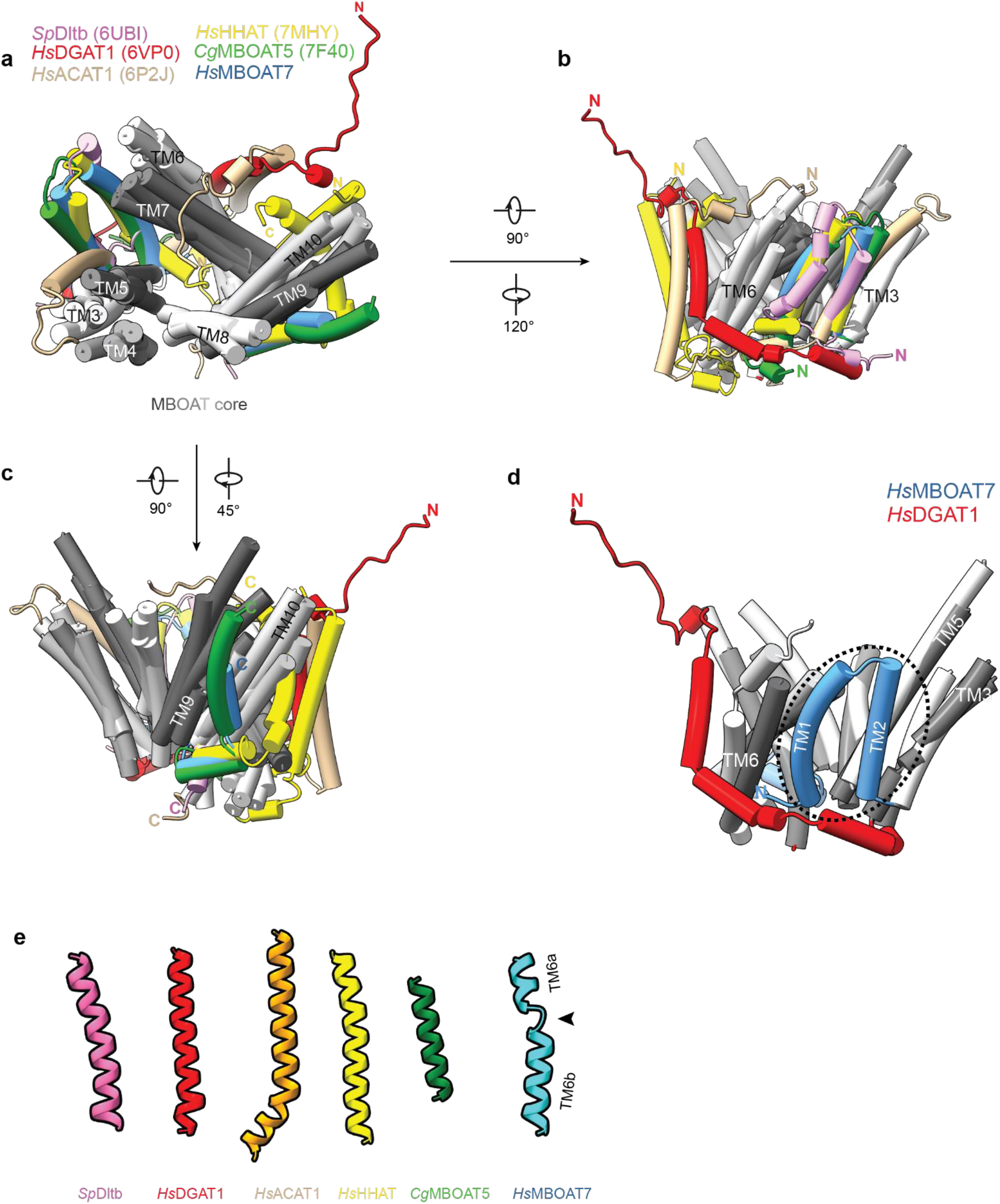
Comparison of MBOAT structures. **a**, MBOAT structures are superimposed. The TMs of the MBOAT7 core are shown in different shades of grey, and the distinct N- and C-termini are colored so as to be consistent with the legend colors. **b** and **c**, Two different orientations to present the variable N (b) and C (c) terminal regions. **d**, Comparison between MBOAT7 and DGAT1. MBOAT7’s N terminal TM1-TM2 (cyan) blocks the lateral gate, and DGAT1’s N-terminus (red) is longer and involved in protein dimerization. **e**, Comparison of MBOAT7 TM6 to the corresponding TMs of other MBOAT proteins. A black arrow denotes the break of MBOAT7 TM6.

**Extended Data Fig. 4.**
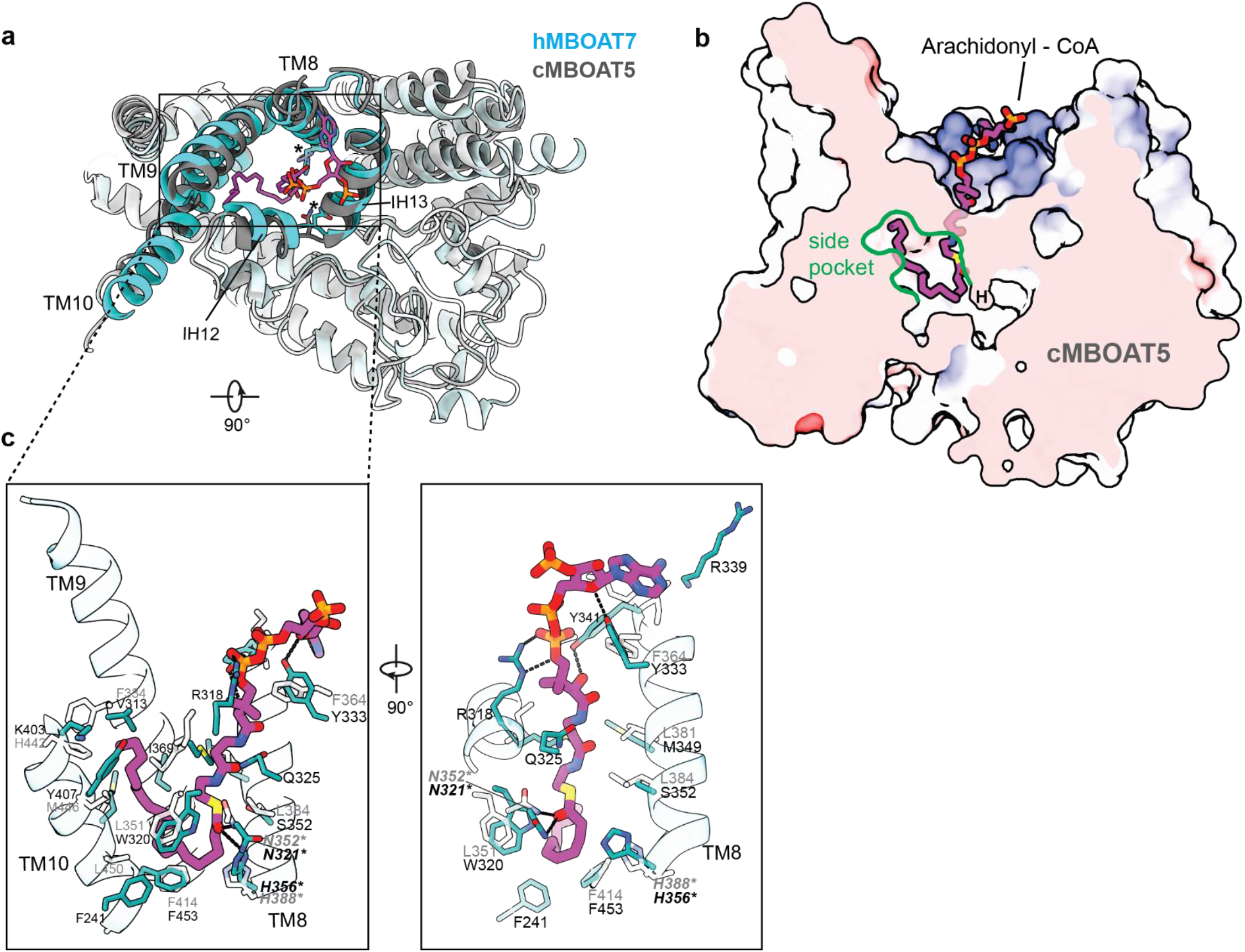
Comparison of the acyl-CoA access channels in human MBOAT7 and chicken MBOAT5. **a**, Top view of the arachidonyl-CoA molecule docked into the MBOAT7 cytosolic channel. Human MBOAT7 (hMBOAT7) structure is superimposed with the chicken MBOAT5 structure complexed with arachidonyl-CoA (PDB 7F40). **b**, A cutting-in view of the cMBOAT5 catalytic chamber and side pocket bound with arachidonyl-CoA, for comparison with Fig. 2b. MBOAT5 is represented with electrostatic surfaces. H indicates the catalytic histidine. **c**, The interaction between arachidonyl-CoA and MBOAT7 residues. The view is the same to Fig. 2c except that the corresponding residues from cMBOAT5 (grey) are labeled in pair.

**Extended Data Fig. 5.**
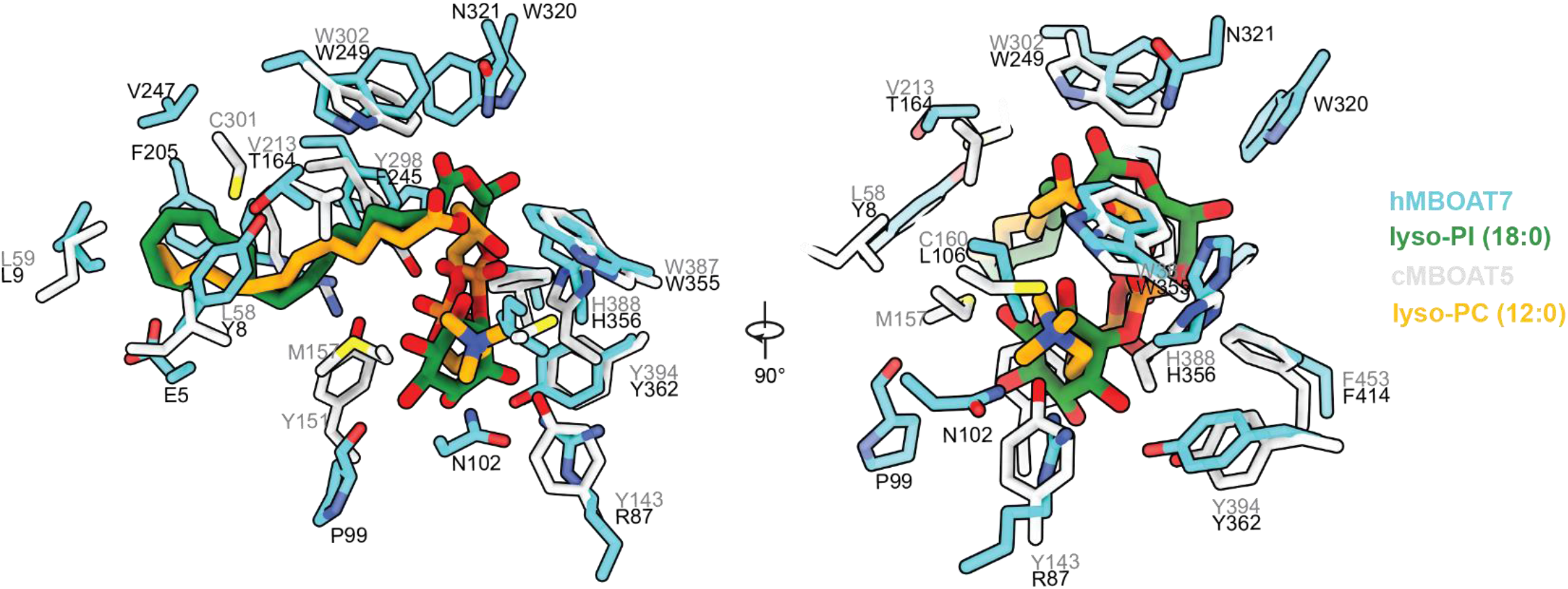
Comparison of the lyso-phospholipid interaction patterns in human MBOAT7 and chicken MBOAT5. The cMBOAT5 with lyso-PC structure (PDB 7F3X) is superimposed with hMBOAT7 with lyso-PI structure. Only the lyso-phospholipid molecules and the key residues are shown.

**Extended Data Fig. 6.**
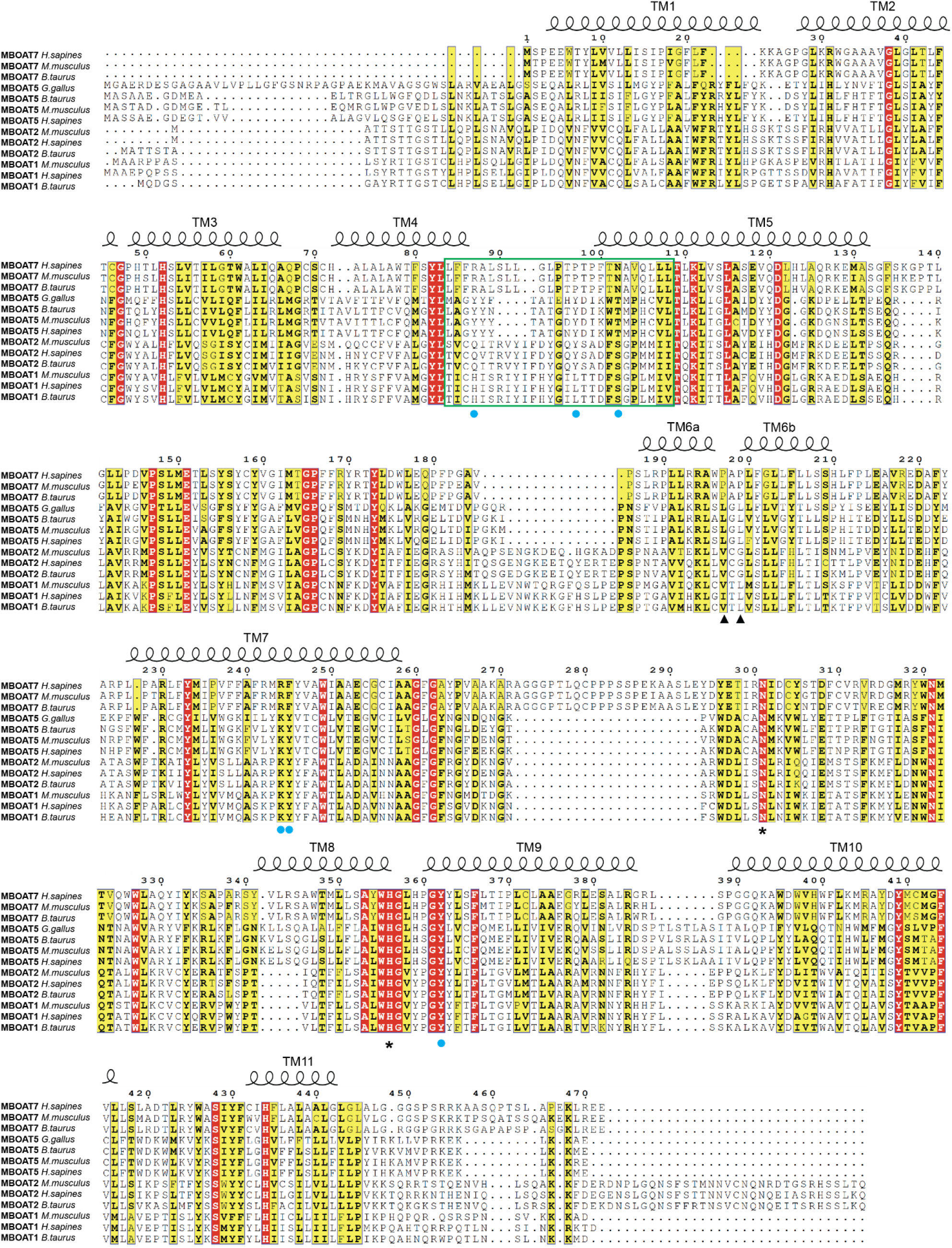
Sequence alignments of MBOAT1, 2, 5 and 7. The sequences were aligned using the Clustal Omega Server. Secondary structural elements of human MBOAT7 are marked above the alignment. Residues were colored based on their conservation using the ESPript server. Catalytic residues are denoted by asterisks. Two conserved proline residues of MBOAT7 were denoted by solid triangles. Important residues that interact with the inositol groups are denoted by solid cyan dots. The variable regions that determine the substrate specificity is highlighted by a green frame.

**Extended Data Fig. 7.**
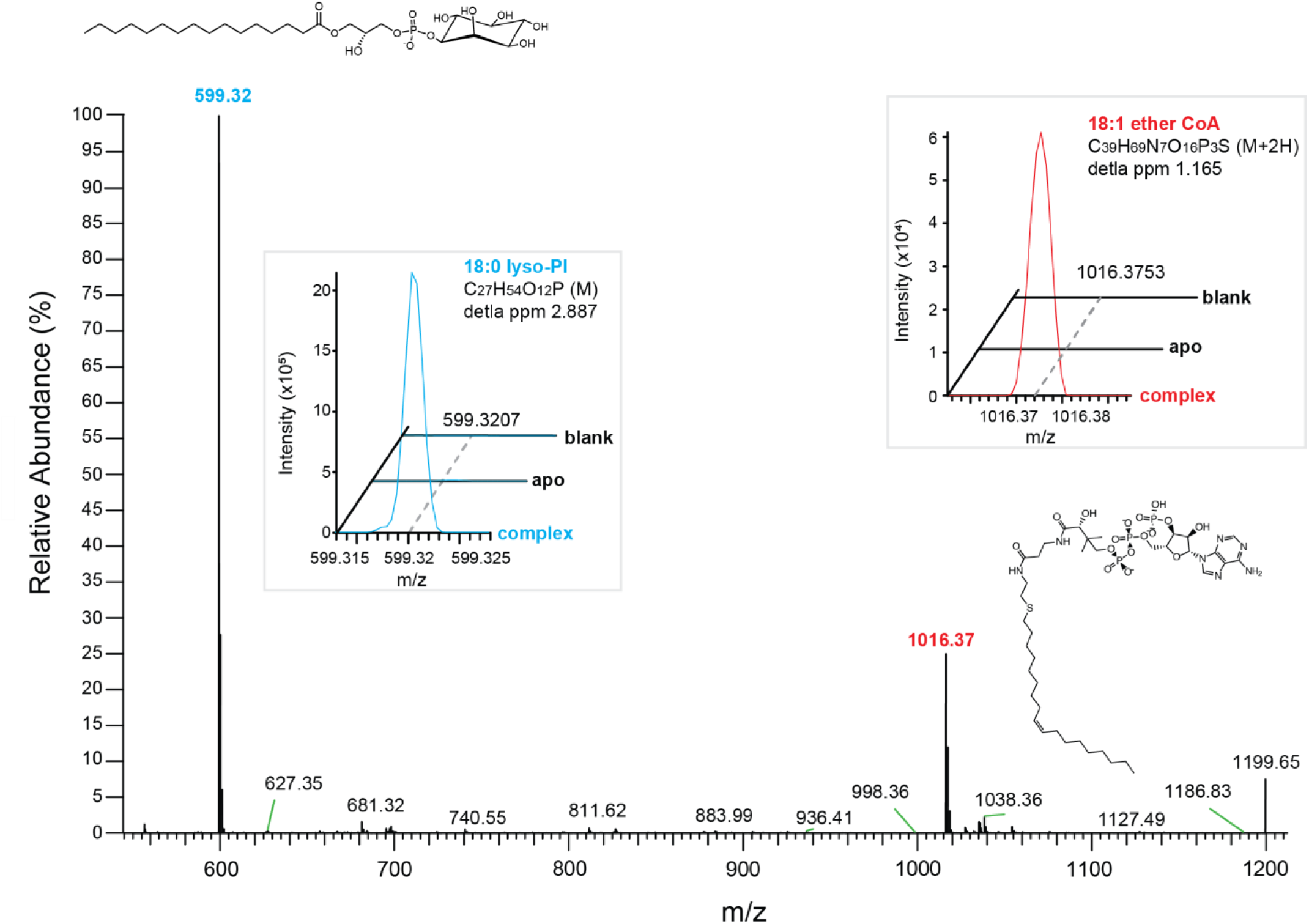
Mass spectrometry analysis of the substrate incorporation into the complex protein prep. The main panel shows the spectrum of the standards of two substrates. Their m/z values and molecular structures are shown as well. Two inserts show the quantification of individual substrates, 18:0 lyso-PI (cyan) and 18:1 ether CoA (red) in blank, apo-protein prep and the complex protein prep.

**Extended Data Fig. 8.**
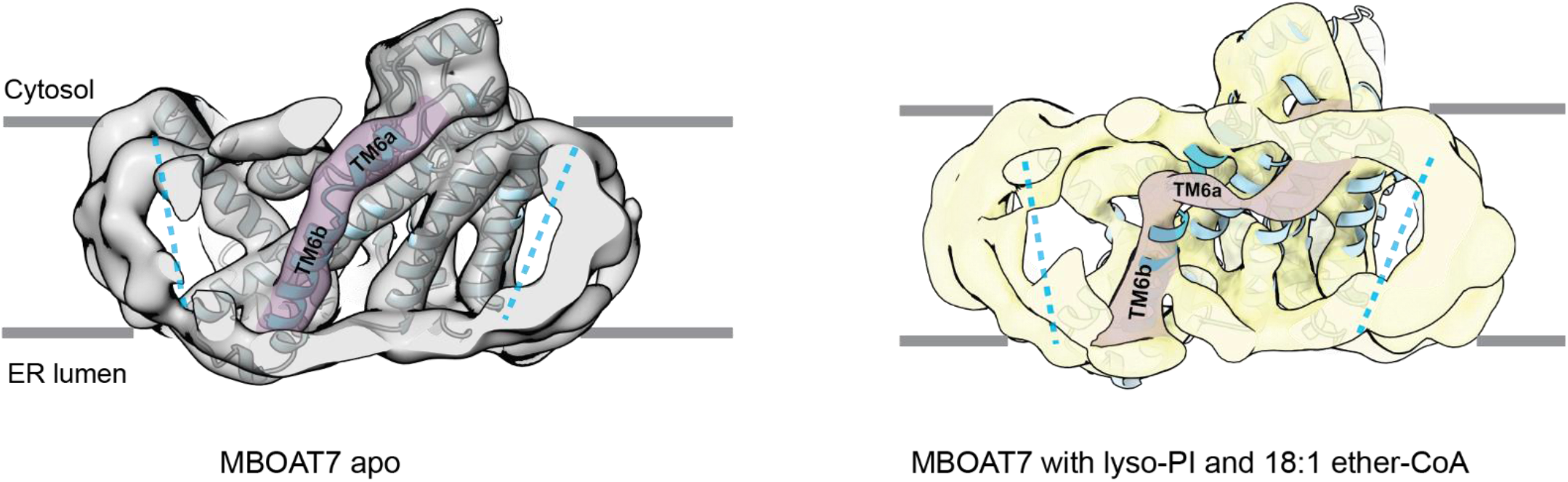
Comparison of MBOAT7 apo-structure and MBOAT7 structure with lyso-PI and 18:1 ether-CoA. The conformational change of TM6 is highlighted in purple. MBOAT7 apo map is low-passed to 6Å, and both maps are contoured at 6 σ. MBOAT7 apo-model is docked into both maps.

**Extended Data Fig. 9.**
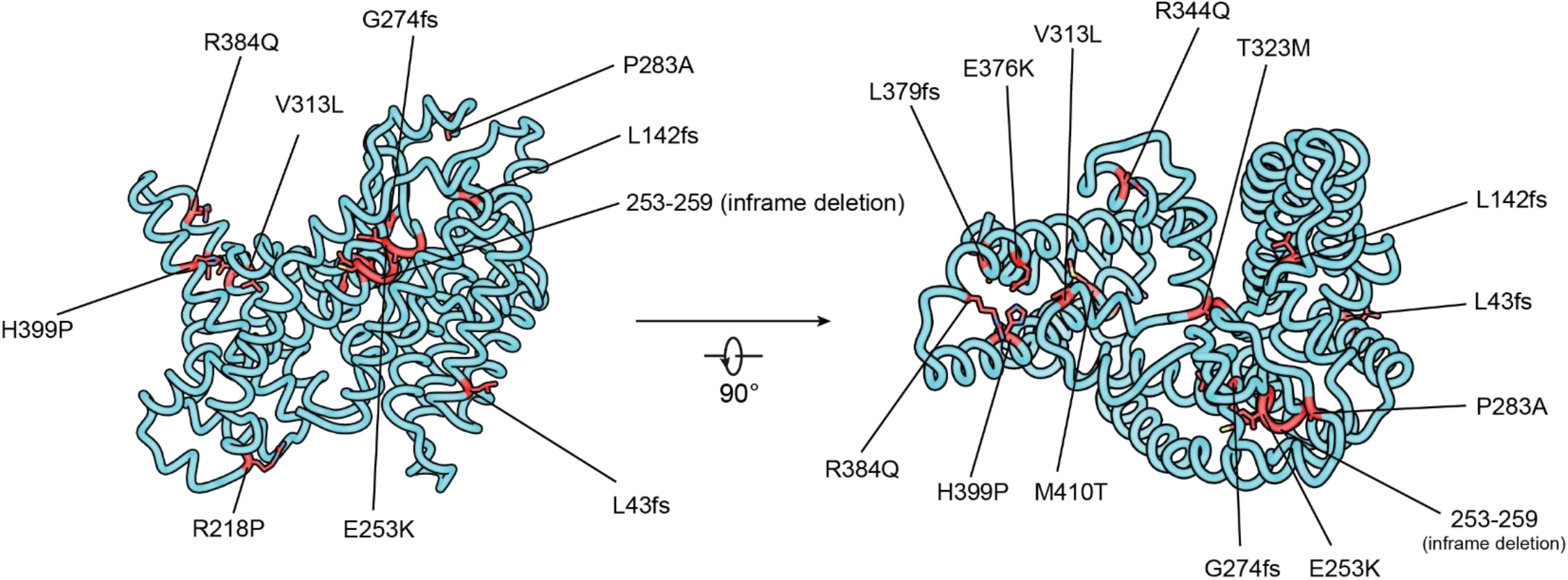
Mapping disease-causing mutations to the MBOAT7 structure. The disease-causing mutations are highlighted in red with side chains. These mutations are correlated with forms of intellectual developmental disorders. fs, frame shift.

## References

1. Vance, J.E. Phospholipid Synthesis and Transport in Mammalian Cells. Traffic 16, 1–18 (2015).

2. Kennedy, E.P. & Weiss, S.B. The function of cytidine coenzymes in the biosynthesis of phospholipides. J Biol Chem 222, 193–214 (1956).

3. Gimeno, R.E. & Cao, J. Thematic Review Series: Glycerolipids. Mammalian glycerol-3-phosphate acyltransferases: new genes for an old activity. Journal of Lipid Research 49, 2079–2088 (2008).

4. Lands, W.E. Metabolism of glycerolipides; a comparison of lecithin and triglyceride synthesis. J Biol Chem 231, 883–8 (1958).

5. Shindou, H. & Shimizu, T. Acyl-CoA:lysophospholipid acyltransferases. J Biol Chem 284, 1–5 (2009).

6. Zhao, Y. et al. Identification and characterization of a major liver lysophosphatidylcholine acyltransferase. J Biol Chem 283, 8258–65 (2008).

7. Gijon, M.A., Riekhof, W.R., Zarini, S., Murphy, R.C. & Voelker, D.R. Lysophospholipid acyltransferases and arachidonate recycling in human neutrophils. J Biol Chem 283, 30235–45 (2008).

8. Catherine C. Y. Chang, J.S., Ta-Yuan Chang. Membrane-bound O-acyltransferases (MBOATs). Front. Biol (2011).

9. Zhang, Q. et al. The structural basis for the phospholipid remodeling by lysophosphatidylcholine acyltransferase 3. Nature Communications 12 (2021).

10. Jiang, Y., Benz, T.L. & Long, S.B. Substrate and product complexes reveal mechanisms of Hedgehog acylation by HHAT. Science 372, 1215–1219 (2021).

11. Coupland, C.E. et al. Structure, mechanism, and inhibition of Hedgehog acyltransferase. Molecular Cell 81, 5025–5038.e10 (2021).

12. Wang, L. et al. Structure and mechanism of human diacylglycerol O-acyltransferase 1. Nature (2020).

13. Sui, X. et al. Structure and catalytic mechanism of a human triacylglycerol-synthesis enzyme. Nature (2020).

14. Qian, H. et al. Structural basis for catalysis and substrate specificity of human ACAT1. Nature (2020).

15. Long, T., Sun, Y., Hassan, A., Qi, X. & Li, X. Structure of nevanimibe-bound tetrameric human ACAT1. Nature (2020).

16. Guan, C. et al. Structural insights into the inhibition mechanism of human sterol O-acyltransferase 1 by a competitive inhibitor. Nat Commun 11, 2478 (2020).

17. Ma, D. et al. Crystal structure of a membrane-bound O-acyltransferase. Nature 562, 286–290 (2018).

18. Liu, Y. et al. Mechanisms and inhibition of Porcupine-mediated Wnt acylation. Nature (2022).

19. Lee, J. et al. Functional and Structural Changes in the Membrane-Bound O-Acyltransferase Family Member 7 (MBOAT7) Protein: The Pathomechanism of a Novel MBOAT7 Variant in Patients With Intellectual Disability. Front Neurol 13, 836954 (2022).

20. Sun, L. et al. Phenotypic Characterization of Intellectual Disability Caused by MBOAT7 Mutation in Two Consanguineous Pakistani Families. Front Pediatr 8, 585053 (2020).

21. Johansen, A. et al. Mutations in MBOAT7, Encoding Lysophosphatidylinositol Acyltransferase I, Lead to Intellectual Disability Accompanied by Epilepsy and Autistic Features. The American Journal of Human Genetics 99, 912–916 (2016).

22. Helsley, R.N. et al. Obesity-linked suppression of membrane-bound O-acyltransferase 7 (MBOAT7) drives non-alcoholic fatty liver disease. Elife 8 (2019).

23. Thabet, K. et al. The membrane-bound O-acyltransferase domain-containing 7 variant rs641738 increases inflammation and fibrosis in chronic hepatitis B. Hepatology 65, 1840–1850 (2017).

24. Thabet, K. et al. MBOAT7 rs641738 increases risk of liver inflammation and transition to fibrosis in chronic hepatitis C. Nat Commun 7, 12757 (2016).

25. Mancina, R.M. et al. The MBOAT7-TMC4 Variant rs641738 Increases Risk of Nonalcoholic Fatty Liver Disease in Individuals of European Descent. Gastroenterology 150, 1219–1230 e6 (2016).

26. Luukkonen, P.K. et al. The MBOAT7 variant rs641738 alters hepatic phosphatidylinositols and increases severity of non-alcoholic fatty liver disease in humans. J Hepatol 65, 1263–1265 (2016).

27. Buch, S. et al. A genome-wide association study confirms PNPLA3 and identifies TM6SF2 and MBOAT7 as risk loci for alcohol-related cirrhosis. Nat Genet 47, 1443–8 (2015).

28. Neumann, C.K.A. et al. MBOAT7-driven phosphatidylinositol remodeling promotes the progression of clear cell renal carcinoma. Mol Metab 34, 136–145 (2020).

29. Donati, B. et al. MBOAT7 rs641738 variant and hepatocellular carcinoma in non-cirrhotic individuals. Scientific Reports 7 (2017).

30. Lu, X., Lin, S., Chang, C.C.Y. & Chang, T.-Y. Mutant Acyl-coenzyme A:Cholesterol Acyltransferase 1 Devoid of Cysteine Residues Remains Catalytically Active. Journal of Biological Chemistry 277, 711–718 (2002).

31. Tunyasuvunakool, K. et al. Highly accurate protein structure prediction for the human proteome. Nature (2021).

32. Jumper, J. et al. Highly accurate protein structure prediction with AlphaFold. Nature (2021).

33. Gorgulla, C. et al. An open-source drug discovery platform enables ultra-large virtual screens. Nature 580, 663–668 (2020).

34. LaPensee, C.R. et al. ATR-101, a Selective and Potent Inhibitor of Acyl-CoA Acyltransferase 1, Induces Apoptosis in H295R Adrenocortical Cells and in the Adrenal Cortex of Dogs. Endocrinology 157, 1775–88 (2016).

35. You, L. et al. Development of a triazole class of highly potent Porcn inhibitors. Bioorganic & Medicinal Chemistry Letters 26, 5891–5895 (2016).

36. Reed, A. et al. LPCAT3 Inhibitors Remodel the Polyunsaturated Phospholipid Content of Human Cells and Protect from Ferroptosis. ACS Chem Biol 17, 1607–1618 (2022).

37. Kowarz, E., Loscher, D. & Marschalek, R. Optimized Sleeping Beauty transposons rapidly generate stable transgenic cell lines. Biotechnol J 10, 647–53 (2015).

38. Zheng, S.Q. et al. MotionCor2: anisotropic correction of beam-induced motion for improved cryo-electron microscopy. Nat Methods 14, 331–332 (2017).

39. Rohou, A. & Grigorieff, N. CTFFIND4: Fast and accurate defocus estimation from electron micrographs. J Struct Biol 192, 216–21 (2015).

40. Thelot, F.A. et al. Distinct allosteric mechanisms of first-generation MsbA inhibitors. Science 374, 580–585 (2021).

41. Bepler, T. et al. Positive-unlabeled convolutional neural networks for particle picking in cryo-electron micrographs. Nat Methods 16, 1153–1160 (2019).

42. Punjani, A., Rubinstein, J.L., Fleet, D.J. & Brubaker, M.A. cryoSPARC: algorithms for rapid unsupervised cryo-EM structure determination. Nature Methods 14, 290–296 (2017).

43. Scheres, S.H. RELION: implementation of a Bayesian approach to cryo-EM structure determination. J Struct Biol 180, 519–30 (2012).

44. Adams, P.D. et al. PHENIX: a comprehensive Python-based system for macromolecular structure solution. Acta Crystallogr D Biol Crystallogr 66, 213–21 (2010).

45. Emsley, P. & Cowtan, K. Coot: model-building tools for molecular graphics. Acta Crystallogr D Biol Crystallogr 60, 2126–32 (2004).

46. Chen, V.B. et al. MolProbity: all-atom structure validation for macromolecular crystallography. Acta Crystallogr D Biol Crystallogr 66, 12–21 (2010).

47. Matyash, V., Liebisch, G., Kurzchalia, T.V., Shevchenko, A. & Schwudke, D. Lipid extraction by methyl-tert-butyl ether for high-throughput lipidomics. J Lipid Res 49, 1137–46 (2008).

48. Van Der Spoel, D. et al. GROMACS: Fast, flexible, and free. Journal of Computational Chemistry 26, 1701–1718 (2005).

49. Klauda, J.B. et al. Update of the CHARMM All-Atom Additive Force Field for Lipids: Validation on Six Lipid Types. The Journal of Physical Chemistry B 114, 7830–7843 (2010).

50. Best, R.B. et al. Optimization of the Additive CHARMM All-Atom Protein Force Field Targeting Improved Sampling of the Backbone φ, ψ and Side-Chain χ1 and χ2 Dihedral Angles. Journal of Chemical Theory and Computation 8, 3257–3273 (2012).

51. Essmann, U. et al. A smooth particle mesh Ewald method. The Journal of Chemical Physics 103, 8577–8593 (1995).

52. Hess, B. P-LINCS: A Parallel Linear Constraint Solver for Molecular Simulation. Journal of Chemical Theory and Computation 4, 116–122 (2008).

53. Nosé, S. A unified formulation of the constant temperature molecular dynamics methods. The Journal of Chemical Physics 81, 511–519 (1984).

54. Hoover, W.G. Canonical dynamics: Equilibrium phase-space distributions. Physical Review A 31, 1695–1697 (1985).

55. Parrinello, M. & Rahman, A. Polymorphic transitions in single crystals: A new molecular dynamics method. Journal of Applied Physics 52, 7182–7190 (1981).

56. Michaud-Agrawal, N., Denning, E.J., Woolf, T.B. & Beckstein, O. MDAnalysis: A toolkit for the analysis of molecular dynamics simulations. Journal of Computational Chemistry 32, 2319–2327 (2011).

57. Wang, J. & Dokholyan, N.V. MedusaDock 2.0: Efficient and Accurate Protein–Ligand Docking With Constraints. Journal of Chemical Information and Modeling 59, 2509–2515 (2019).

58. Morris, G.M. et al. AutoDock4 and AutoDockTools4: Automated docking with selective receptor flexibility. J Comput Chem 30, 2785–91 (2009).

